# DeepCob: Precise and high-throughput analysis of maize cob geometry using deep learning with an application in genebank phenomics

**DOI:** 10.1101/2021.03.16.435660

**Authors:** Lydia Kienbaum, Miguel Correa Abondano, Raul Blas, Karl Schmid

## Abstract

**Background:** Maize cobs are an important component of crop yield that exhibit a high diversity in size, shape and color in native landraces and modern varieties. Various phenotyping approaches were developed to measure maize cob parameters in a high throughput fashion. More recently, deep learning methods like convolutional neural networks (CNN) became available and were shown to be highly useful for high-throughput plant phenotyping. We aimed at comparing classical image segmentation with deep learning methods for maize cob image segmentation and phenotyping using a large image dataset of native maize landrace diversity from Peru.

**Results:** Comparison of three image analysis methods showed that a Mask R-CNN trained on a diverse set of maize cob images was highly superior to classical image analysis using the Felzenszwalb algorithm and a Window-based CNN due to its robustness to image quality and object segmentation accuracy (*r* = 0.99). We integrated Mask R-CNN into a high-throughput pipeline to segment both maize cobs and rulers in images and perform an automated quantitative analysis of eight phenotypic traits, including diameter, length, ellipticity, asymmetry, aspect ratio and average RGB values for cob color. Statistical analysis identified key training parameters for efficient iterative model updating. We also show that a small number of 10-20 images is sufficient to update the initial Mask R-CNN model to process new types of cob images. To demonstrate an application of the pipeline we analyzed phenotypic variation in 19,867 maize cobs extracted from 3,449 images of 2,484 accessions from the maize genebank of Peru to identify phenotypically homogeneous and heterogeneous genebank accessions using multivariate clustering.

**Conclusions:** Single Mask R-CNN model and associated analysis pipeline are widely applicable tools for maize cob phenotyping in contexts like genebank phenomics or plant breeding.

## Background

High-throughput precision phenotyping of plant traits is rapidly becoming an integral part of plant research, plant breeding, and crop production [1]. This development complements the rapid advances in genomic methods that, when combined with phenotyping, enable rapid, accurate, and efficient analysis of plant traits and the interaction of plants with their environment [2]. However, for many traits of interest, plant phenotyping is still labor intensive or technically challenging. Such a bottleneck in phenotyping [3] limits progress in understanding the relationship between genotype and phenotype, which is a problem for plant breeding [4]. The phenotyping bottleneck is being addressed by phenomics platforms that integrate high-throughput automated phenotyping with analysis software to obtain accurate measurements of phenotypic traits [5, 6]. Existing phenomics platforms cover multiple spatial and temporal scales and incorporate technologies such as RGB image analysis, NIRS, or NMR spectroscopy [7, 8, 9]. The rapid and large-scale generation of diverse phenotypic data requires automated analysis to convert the output of phenotyping platforms into meaningful information such as measures of biological quantities [10, 11]. Thus, high-throughput pipelines with accurate computational analysis will realize the potential of plant phenomics by overcoming the phenotyping bottleneck.

A widely used method for plant phenotyping is image segmentation and shape analysis using geometric morphometrics [12]. Images are captured in standardized environments and then analyzed either manually or automatically using image annotation methods to segment images and label objects. The key challenge in automated image analysis is the detection and segmentation of relevant objects. Traditionally, object detection in computer vision (CV) has been performed using multi-variate algorithms that detect edges, for example. Most existing pipelines using classical image analysis in plant phenotyping are species-dependent and assume homogeneous plant material and standardized images [13, 14, 15]. Another disadvantage of classical image analysis methods is low accuracy and specificity when image quality is low or background noise is present. Therefore, the optimal parameters for image segmentation often need to be fine-tuned manually through experimentation. In recent years, machine learning approaches have revolutionized many areas of CV such as object recognition [16] and are superior to classical CV methods in many applications [17]. The success of machine learning in image analysis can be attributed to the evolution of neural networks from simple architectures to advanced feature-extracting convolutional neural networks (CNN) [18]. The complexity of CNN could be exploited because deep learning algorithms offered new and improved training approaches for these more complex method networks. Another advantage of machine learning methods is their robustness to variable image backgrounds and image qualities when model training is based on a sufficiently diverse set of training images. Although CNN have been very successful in general image classification and segmentation, their application in plant phenotyping is still limited to a few species and features. Current applications include plant pathogen detection, organ and feature quantification, and phenological analysis [19, 20, 9].

Maize cobs can be described with few geometric shape and color parameters. Since the size and shape of maize cobs are important yield components with a high heritability and are correlated with total yield [21, 22], they are potentially useful traits for selection in breeding programs. High throughput phenotyping approaches are also useful for characterizing native diversity of crop plants to facilitate their conservation or utilize them as genetic resources [23, 24]. Maize is an excellent example to demonstrate the usefulness of high throughput phenotyping because of its high genetic and phenotypic diversity, which originated since its domestication in South-Central Mexico about 9,000 years ago [25, 26, 27]. A high environmental variation within its cultivation range in combination with artificial selection by humans resulted in many phenotypically divergent landraces [28, 29]. Since maize is one of the most important crops worldwide, large collections of its native diversity were established in *ex situ* genebanks, whose genetic and phenotypic diversity are now being characterized [30]. This unique pool of genetic and phenotypic variation is threatened by genetic erosion [31, 32, 33] and understanding its role in environmental and agronomic adaptation is essential to identify valuable genetic resources and develop targeted conservation strategies.

In the context of native maize diversity we present a CNN-based deep learning model implemented in a robust and widely applicable analysis pipeline for recognizing, semantic labeling and automated measurements of maize cobs in RGB images for large scale plant phenotyping. Highly variable traits like cob length, kernel color and number were used for classification of the native maize diversity of Peru [34] and are useful for the characterization of maize genetic resources because cobs are easily stored and field collections can be analyzed at a later time point. We demonstrate the application of image segmentation to photographs of native maize diversity in Peru. So far, cob traits have been studied for small sets of Peruvian landraces, only such as cob diameter in 96 accessions of 12 Peruvian maize landraces [35], or cob diameter in 59 accessions of 9 highland landraces [36]. Here we use image analysis to obtain cob parameters from 2,484 accessions of the Peruvian maize genebank hosted at Universidad Nacional Agraria La Molina (UNALM) by automated image analysis. We also show that the DeepCob image analysis pipeline can be easily expanded to different image types of maize cobs such as segregating populations resulting from genetic crosses.

## Results

### Comparison of image segmentation methods

To address large-scale segmentation of maize cobs, we compared three different image analysis methods for their specificity and accuracy in detecting and segmenting both maize cobs and measurement rulers in RGB images. Correlations between true and derived values for cob length and diameter show that Mask R-CNN far outperformed the classical image segmentation method Felzenszwalb (FelzSEG) and a window-based CNN (WindowCNN) (Figure 1). For two sets of old (ImgOld) and new (ImgNew) maize cob images (see Materials and Methods), Mask R-CNN achieved correlations of 0.99 and 1.00, respectively, while correlation coefficients ranged from 0.14 to 0.93 with FelzSEG and from 0.03 to 0.42 with WindowCNN, respectively. Since Mask R-CNN was strongly superior in accuracy to the other two segmentation methods, we restricted all further analyses to this method only.

**Figure 1:**
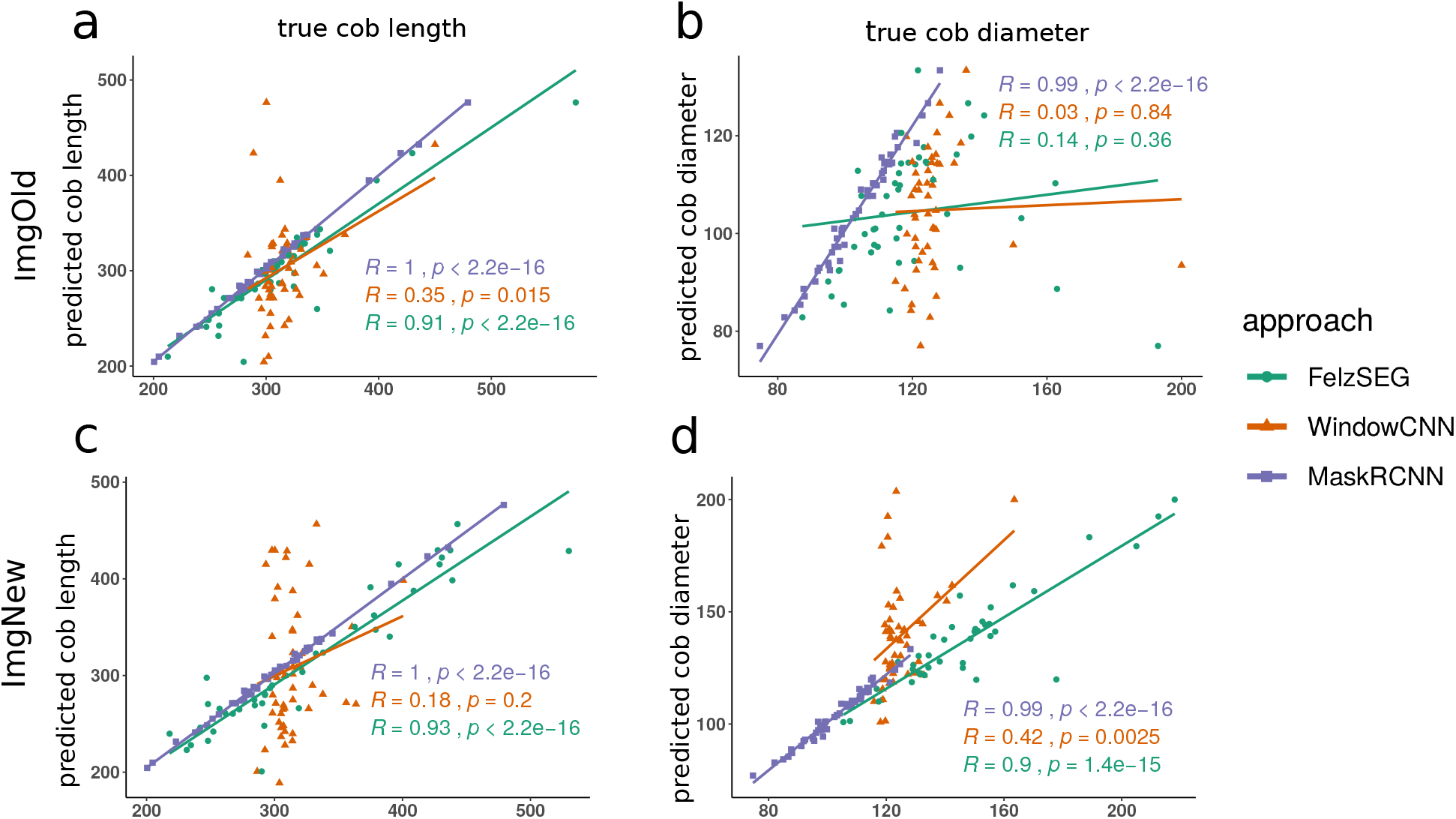
Pearson correlation between true and estimated cob length for three image segmentation methods (FelzSEG,WindowCNN,Mask R-CNN). True (x-axis) and estimated (y-axis) mean cob length (a,c) and diameter (b,d) per image with each approach, split by dataset, ImgOld and ImgNew are shown. In all cases, *MaskRCNN* achieves the highest correlation of at least 0.99 with the true values.

### Parameter optimization of Mask R-CNN

We first describe parameter optimizations during training of the Mask R-CNN model based on the old (ImgOld) and new (ImgNew) maize cob image data from the Peruvian maize genebank. A total of 90 models were trained, differing by the parameters *learning rate, total epochs, epochs*.*m, mask loss weight, monitor, minimask* (see Material and Methods), using a small (200) and a large (1,000) set of randomly selected images as training data. The accuracy of Mask R-CNN detection depends strongly on model parameters, as *AP*@[.5:.95] values for all models ranged from 5.57 to 86.74 for 200 images and from 10.49 to 84.31 for 1,000 images for model training (Supplementary Table S1). Among all 90 models, M104 was the best model for maize cob and ruler segmentation with a score of 86.74, followed by models M101, M107, and M124 with scores of 86.56. All four models were trained with the small image dataset.

Given the high variation of the scores, we evaluated the contribution of each training parameter to this variation with an ANOVA (Table 1). There is an interaction effect between the size of the training set and the total number of epochs trained, as well as an effect of a minimask, which is often used as a resizing step of the object mask before fitting it to the deep learning model. The other training parameters *learning rate, monitoring, epochs*.*m* (mode to train only heads or all layers), and *mask loss weight* had no effect on the *AP*@[.5:.95] value. The lsmeans show that training without *minimask* leads to higher scores and more accurate object detection. Table 1 shows an interaction between the size of the training set and the total number of epochs. Model training with 200 images over 200 epochs was not different from training over 50 epochs or from model training with 1,000 images over 200 epochs at *p <* 0.05. In contrast, model training over 15 epochs only resulted in lower *AP*@[.5:.95] values.

**Table 1:**
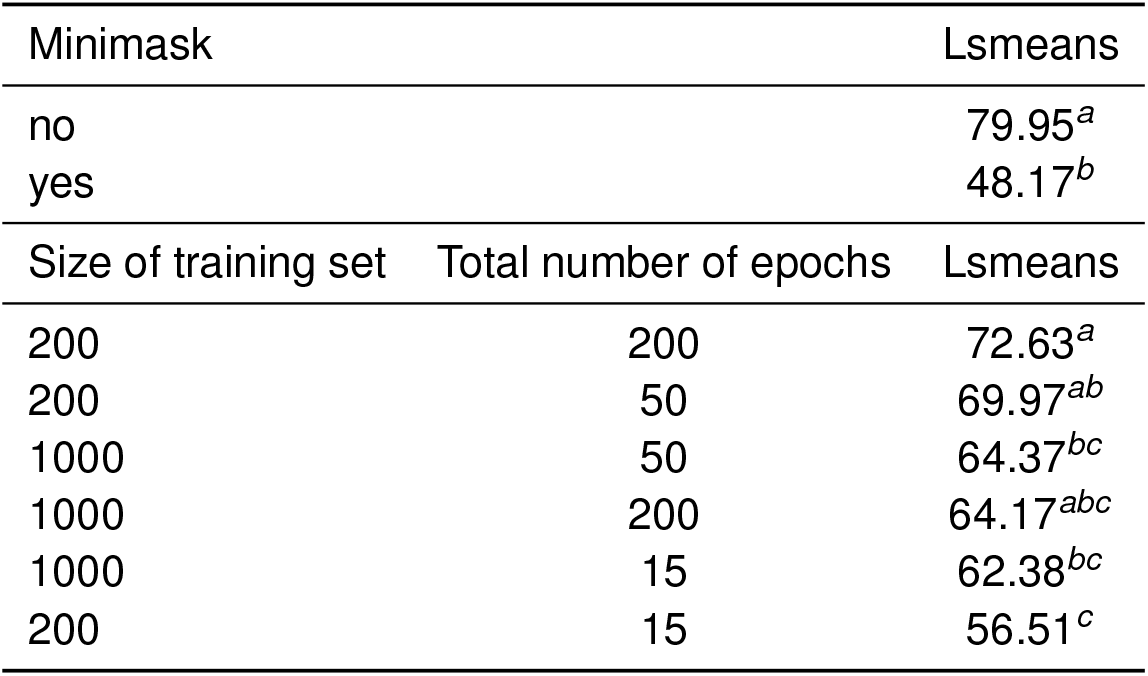
Lsmeans of *AP*@[.5:.95] in the ANOVA analysis for Mask R-CNN model parameters *minimask* and the interaction of *training set size × total number of epochs*. Mean values that share a common letter are not significantly different (*p <* 0.05). Individual *p*-values of comparisons are in Supplementary Tables S2 and S3.

| Minimask |  | Lsmeans |
| --- | --- | --- |
| no |  | 79.95<sup>a</sup> |
| yes |  | 48.17<sup>b</sup> |
| Size of training set | Total number of epochs | Lsmeans |
| 200 | 200 | 72.63<sup>a</sup> |
| 200 | 50 | 69.97<sup>ab</sup> |
| 1000 | 50 | 64.37<sup>bc</sup> |
| 1000 | 200 | 64.17<sup>abc</sup> |
| 1000 | 15 | 62.38<sup>bc</sup> |
| 200 | 15 | 56.51<sup>c</sup> |

### Loss behavior of Mask R-CNN during model training

Monitoring loss functions of model components (classes, masks, boxes) during model training identifes components that need further adjustments to achieve full optimization. Compared to the other components, mask loss contributed the highest proportion to all losses (Figure 2), which indicates that the most challenging process in model training and optimization is segmentation by creating masks for cobs and rulers. The best model M104 shows a decreasing training and validation loss during the first 100 epochs and a tendency for overfitting in additional epochs (Figure 2b). This suggests that model training over 100 epochs is sufficient. Other models like M109 (Figure 2c) exhibit overfitting with a 10-fold higher validation loss than M104. Instead of learning patterns, the model memorizes training data, which increases the validation loss and results in weak predictions for object detection and image segmentation.

**Figure 2:**
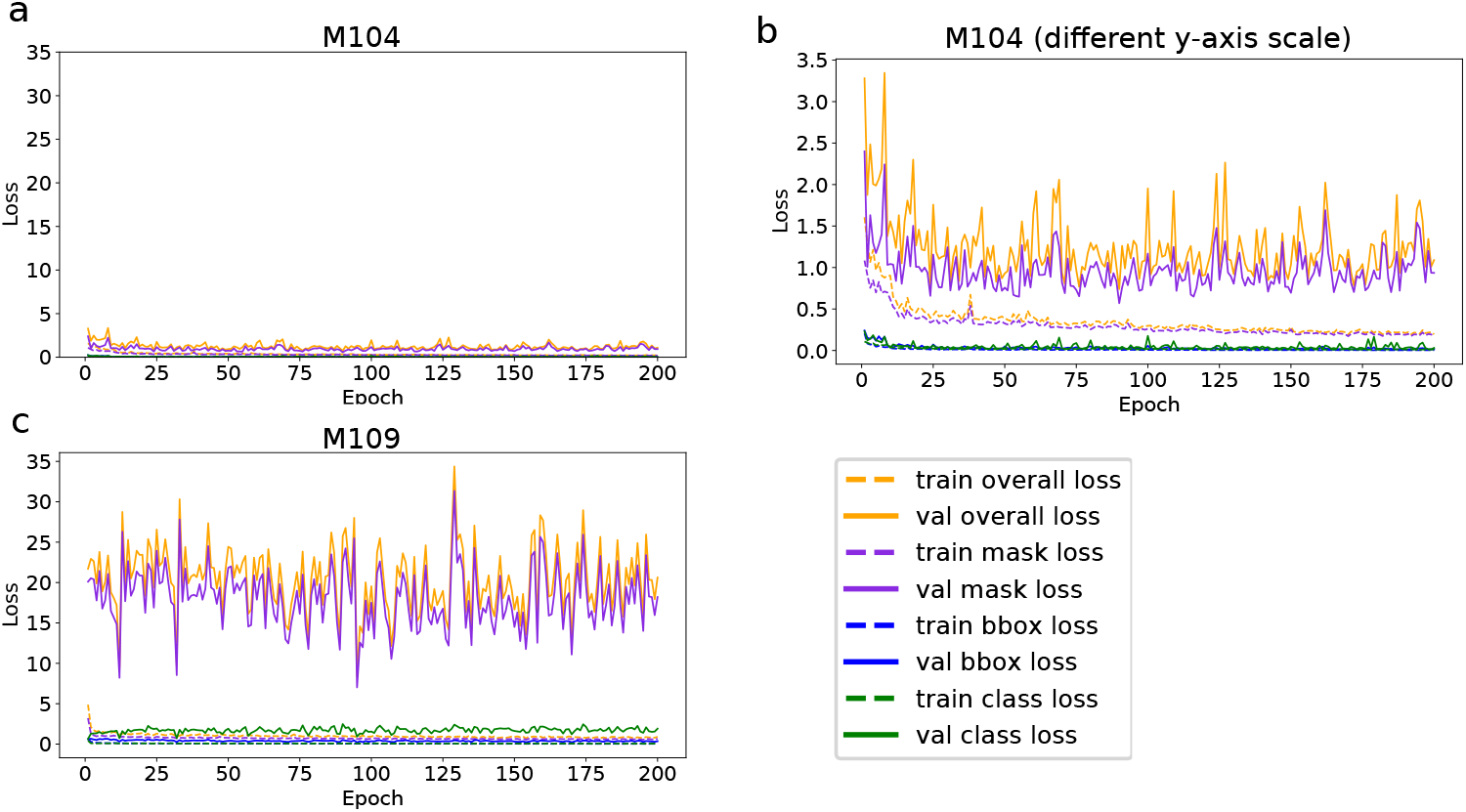
Mask R-CNN training and validation losses during training for 200 epochs on ImgOld and ImgNew maize cob images from the Peruvian genebank. a) Loss curves for model M104, which emerged as the best model b) Model M104 with a different scale on the y-axis. The mask loss showed the largest effect on overall loss, indicating that masks are most difficult to optimize. Other losses, like class loss or bounding box loss, are of minor importance. c) Model M109 shows overfitting as indicated by much higher validation losses resulting in an inferior model based on *AP*@[.5:.95].

### Visualization of feature maps generated by Mask R-CNN

Although neural networks are considered a “black box” method, a feature map visualization of selected layers shows interpretable features of trained networks. In a feature map, high activations correspond to high feature recognition activity in that area, as shown in Figure 3A for the best model M104. Over several successive CNN layers, the cob shape is increasingly well detected until, in the last layer (res4a) the feature map indicates a robust distinction between foreground with the cob and ruler objects and the background. High activations occur at the top of the cobs (Fig. 3A, res4g layer), which may contribute to localization. Because the cobs were oriented according to their lower (apical) end in the images, it may be more difficult for the model to detect the upper edges, which are variable in height. Overall, the feature maps show that the network learned specific features of the maize cob and the image background.

**Figure 3:**
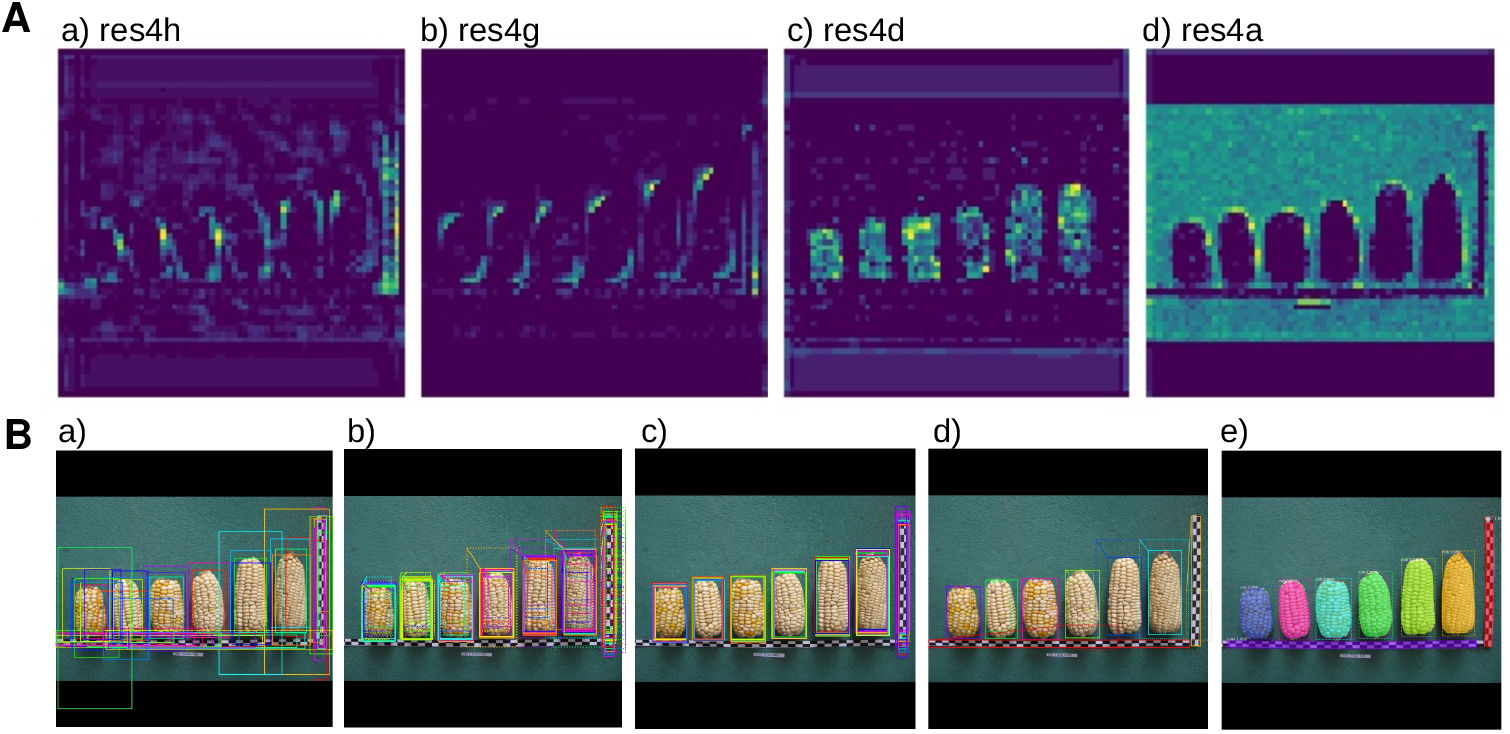
Feature map visualizations and improved segmentation throughout learning A) Examples of feature map visualizations on resnet-101 (for an explanation, see Materials and Methods). a) An early layer shows activations around the cob shape and the ruler on the right. b) The next layer shows more clarified cob shapes with activations mainly at the top and bottom of cobs c) A later layer shows different activations inside the cob. d) The latest layer masks the background very well masked from cobs and rulers. B) Visualization of the main detection procedure of Mask R-CNN a) The top 50 anchors obtained from the region proposal network (RPN), after non-max suppression. b), c) and d) show further bounding box refinement and e) shows the output of the detection network: mask prediction, bounding box prediction and class label. All images are quadratic with a black padding because images are internally resized to a quadratic scale for more efficient matrix multiplication operations.

The Mask R-CNN detection process can be visualized by its main steps, which we demonstrate using the best model (Figure 3B). The top 50 anchors are output by the Region Proposal Network (RPN) and the anchored boxes are then further refined. In the early stages of refinement, all boxes already contain a cob or ruler, but boxes containing the same image element have different lengths and widths. In later stages, the boxes are further reduced in size and refined around the cobs and rulers until, in the final stage, mask recognition provides accurate-fitting masks, bounding boxes, and class labels around each recognized cob and ruler.

The best Mask R-CNN model for detection and segmentation of both maize cobs and rulers is very robust to image quality and variation. This robustness is evident from a representative subset of ImgOld and ImgNew images that we did not use for training and show a high variation in image quality, backgrounds and diversity of maize cobs (Figure 4). Both the identification of bounding boxes and object segmentation are highly accurate regardless of image variability. The only inaccuracies in the location of bounding boxes or masks occur at the bottom edge of cobs.

**Figure 4:**
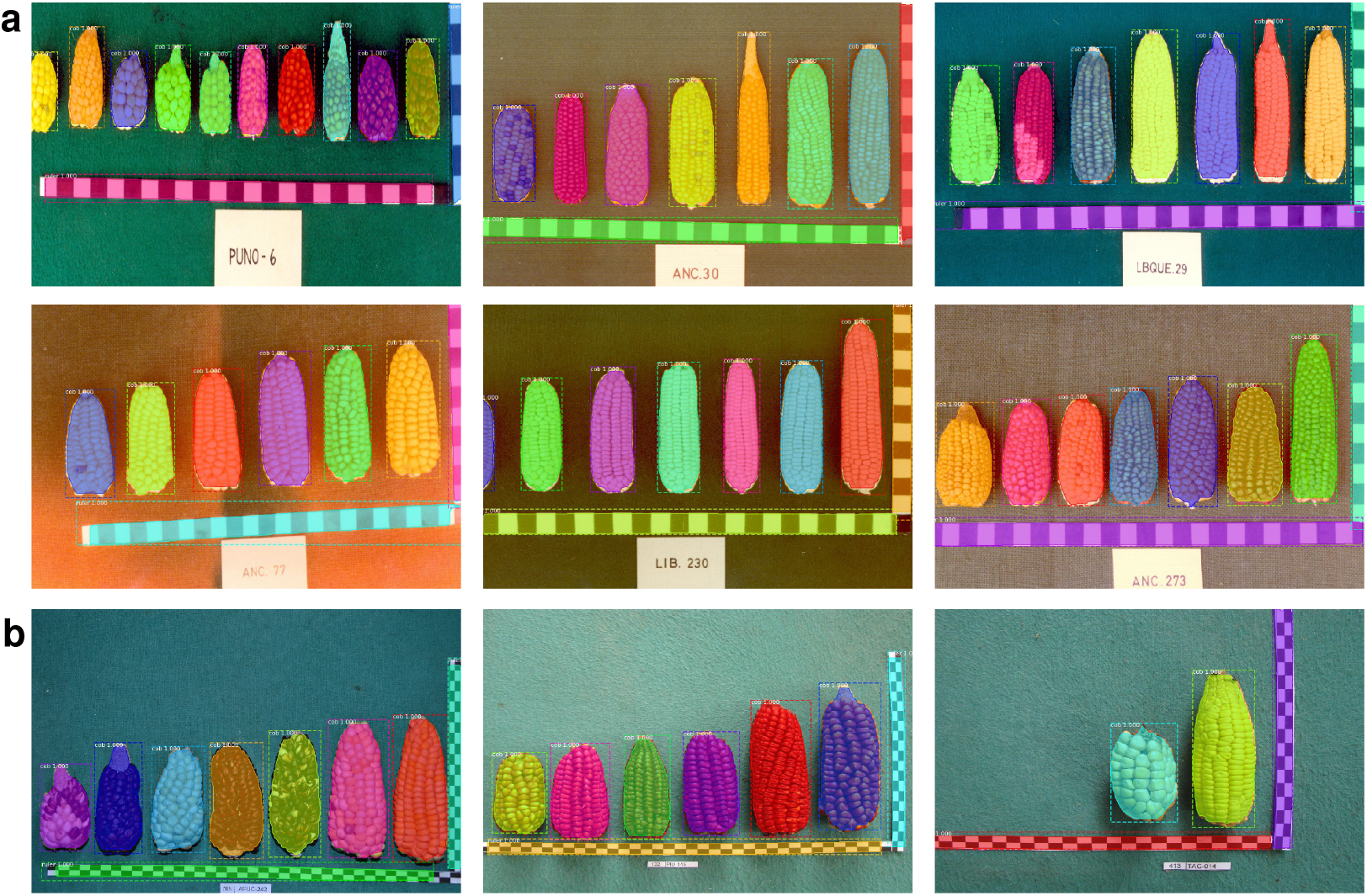
Examples of detection and segmentation performance on a representative example of diverse images from the Peruvian maize landrace ImgOld (a) and ImgNew (b) image sets including different cob and background colors.

### Maize model updating on additional image datasets

To extend the use of our model for images of corn cobs taken under different circumstances and in different environments (e.g., in the field), we investigated whether updating our maize model for new image types with additional image data included in the ImgCross and ImgDiv data sufficiently improves the segmentation accuracy of cob and ruler elements compared to a full training process starting again with the standard COCO model. We used the best maize model trained on ImgOld and ImgNew data (model M104, hereafter maize model), which is pre-trained only on the cob and ruler classes. In addition to updating to our maize model, we updated the COCO model with the same images. In this context, the COCO model serves as a validation, as it is a standard mask-R CNN model trained on the COCO image data [37], which contains 80 annotated object classes in 330K images.

Overall, model updating using training images significantly improved the *AP*@[.5:.95] scores of the additional image datasets (Figure 5), with scores differing between image sets, initial models, and training set sizes. With standard COCO model weights (Fig. 6a, c), *AP*@[.5:.95] scores were initially low, down to a value of 0, in which neither cobs nor rulers were detected. However, scores increased rapidly during up to 0.7 during the first 30 epochs. In contrast, with the pre-trained weights (Fig. 5b, d) of the maize model *AP*@[.5:.95] scores were already high during the first epochs and then rapidly improved to higher values than with the COCO model. Therefore, object segmentation using additional maize cob image data was significantly better with the pre-trained maize model from the beginning and throughout the model update.

**Figure 5:**
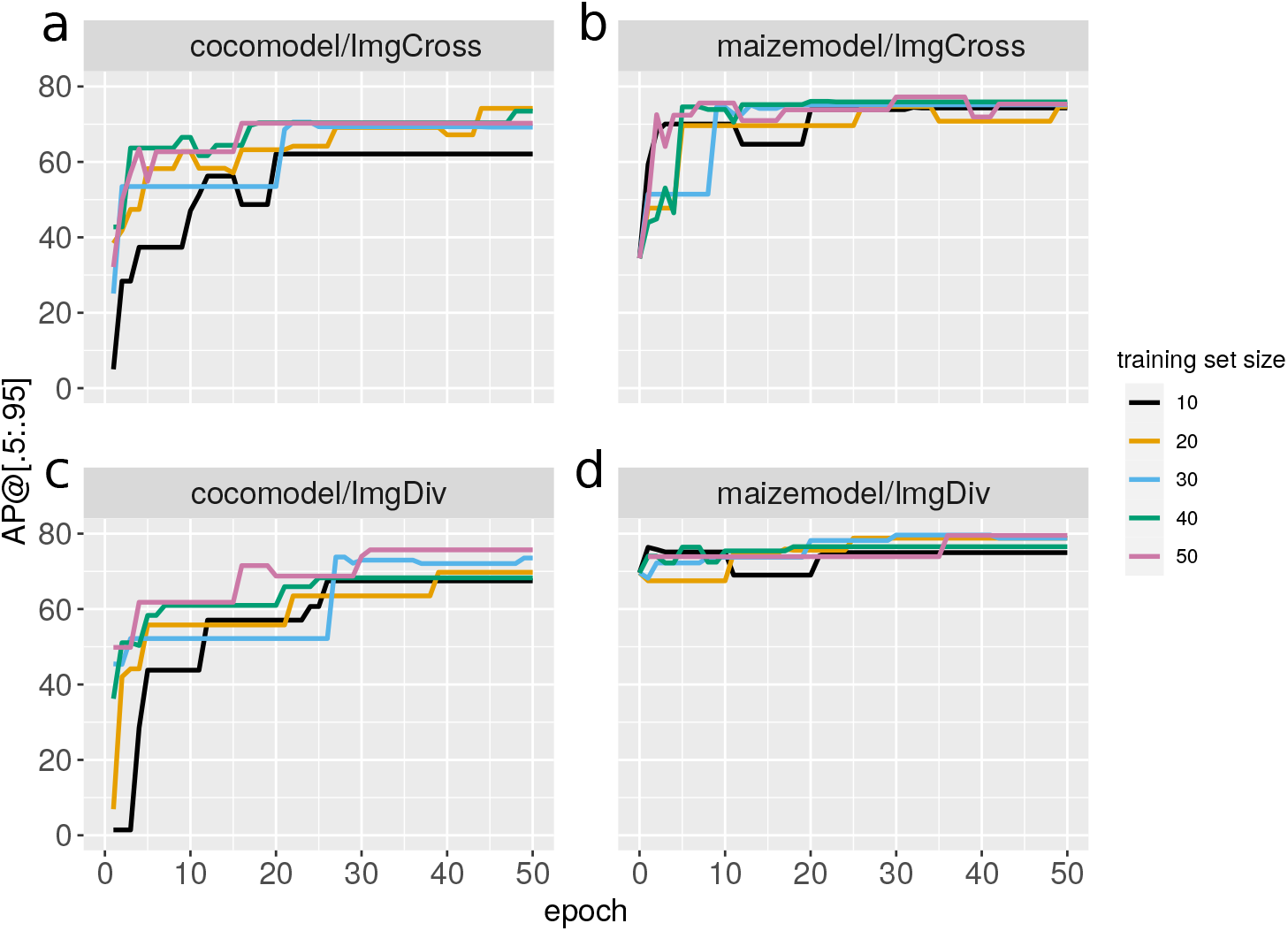
Improvement of *AP*@[.5:.95] scores during 50 epochs of model updating to different maize cob image datasets (a, b: ImgCross; c, d: ImgDiv). Updating on the COCO initial weights/COCO model (a,c) in comparison to updating on the pre-trained maize model (b,d) depends on different amounts of training images, namely 10, 20, 30, 40 or 50 images.

**Figure 6:**
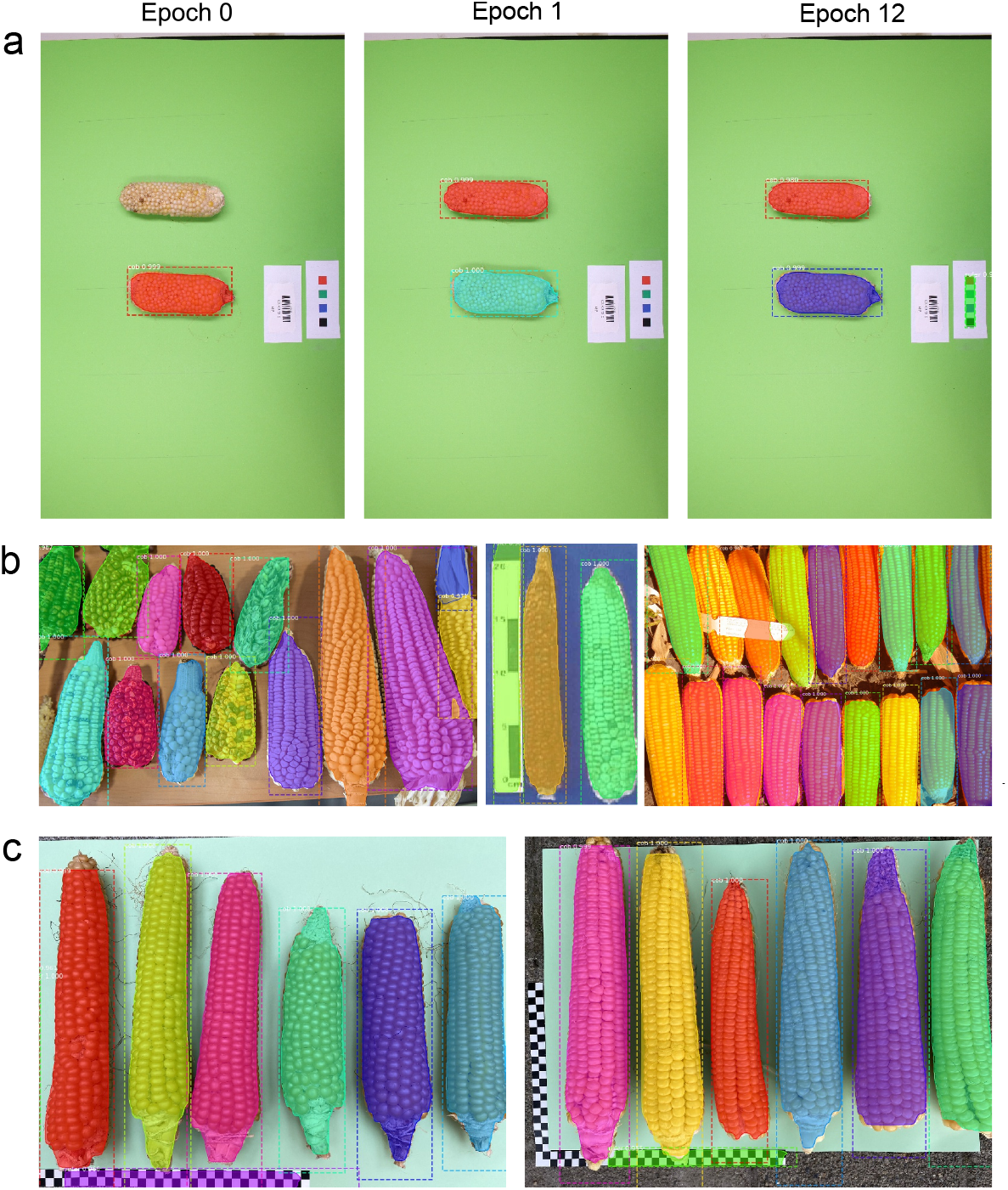
Detection of cob and ruler after model updating the pretained maize model with different image datasets. a) Updating with 10 training images from ImgCross. The original maize model detected only one cob (epoch 0). After one epoch of model updating both cobs were accurately segmented and after epoch 12 the different ruler element was detected. Photo credit: K. Schmid, University of Hohenheim. b) Segmentation of various genebank images after updating for 25 epochs with 20 training images from ImgDiv. Photo credits: https://nexusmedianews.com/ drought-is-crippling-small-farmers-in-mexico-with-consequences-for-everyone-else-photos-73b35a01e4d (Left) https://www.ars.usda.gov/ARSUserFiles/50301000/Races\_of\_Maize/RoM\_Paraguay\_0\_Book.pdf (Center) Right: CIMMYT,https://flic.kr/p/9h9X6B. All photos are available under a Creative Commons License. c) Segmentation of cobs and rulers in post-harvest images of the Swiss Rheintaler Ribelmais landrace with the best model from ImgCross without updating on these images. Photo credit: Benedikt Kogler, Verein Rheintaler Ribelmais e.V., Switzerland

Given the high variation in these scores, we determined the contribution of the three factors *starting model, training set size* and *training data set* to the observed variation in *AP*@[.5:.95] scores with an ANOVA. In this analysis, the interactions between dataset and starting model were significant. By accounting for the lsmeans of these significant interactions (Table 2), updating of the pre-trained maize model than of the COCO model was better in both data sets. With respect to traing set sizes, *AP*@[.5:.95] scores of maize model were essentially the same for different sizes and were always higher than of the COCO model. In summary, there is a clear advantage in updating a pre-trained maize model over the COCO model for cob segmentation with diverse maize cob image sets.

**Table 2:** Lsmeans of *AP*@[.5:.95] score of the significant interactions for model updating, dataset *×* starting model and starting model *×* training set size. Means sharing a common letter are not significantly different.

| Dataset | Starting Model | Lsmeans |
| --- | --- | --- |
| ImgDiv | maize | 75.40<sup>a</sup> |
| ImgCross | maize | 71.04<sup>b</sup> |
| ImgCross | COCO | 62.74<sup>c</sup> |
| ImgDiv | COCO | 61.86<sup>c</sup> |
| Starting Model | Dataset | Lsmeans |
| maize | 40 | 74.11<sup>a</sup> |
| maize | 50 | 74.06<sup>a</sup> |
| maize | 30 | 73.48<sup>a</sup> |
| maize | 10 | 72.40<sup>a</sup> |
| maize | 20 | 72.03<sup>a</sup> |
| COCO | 50 | 67.54<sup>b</sup> |
| COCO | 40 | 65.39<sup>b</sup> |
| COCO | 20 | 61.71<sup>c</sup> |
| COCO | 30 | 61.67<sup>c</sup> |
| COCO | 10 | 55.19<sup>d</sup> |

### Descriptive of data obtained from cob image segmentation

To demonstrate that the Mask R-CNN model is suitable for large-scale and accurate image analysis, we present the results of a descriptive analysis of 19,867 maize cobs that were identified and extracted from the complete set of images from the Peruvian maize genebank, i.e., the ImgOld and ImgNew data. Here, we focus on the question whether image analysis identifies genebank accessions which are highly heterogeneous with respect to cob traits by using measures of trait variation and multivariate clustering algorithms.

Our goal was to identify heterogeneous genebank accessions that either harbor a high level of genetic variation or are admixed because of co-cultivation of different landraces on farmers fields or mix-ups during genebank storage. We therefore analysed variation of cob parameters within images to identify genebank accessions with a high phenotypic diversity of cobs using two different multivariate analysis methods to test the robustness of the classification.

The first approach consisted of calculating a *Z* -score of each cob in an image as measure of deviation from the mean of the image (Within image *Z* -scores), clustering these scores with a PCA, followed by applying CLARA and determining the optimal number of clusters with the average silhouette method. The second approach consisted of calculating a centered and scaled standard deviation of cob parameters for each image, applying a PCA to the values of all images, clustering with *k* -means and determining the optimal cluster number with the gap statistic. With both approaches, the best-fitting numbers of clusters was *k* = 2 with a clear separation between clusters and little overlap along the first principal component (Figure 7). The distribution of trait values between the two groups shows that they differ mainly by the three RGB colors and cob length (in the *Z* -score analysis only) suggesting that cob color tends to more variable than most morphological traits within genebank accessions. Supplementary Figure S1 shows images of genebank accessions classified as homogeneous and variable, respectively.

**Figure 7:**
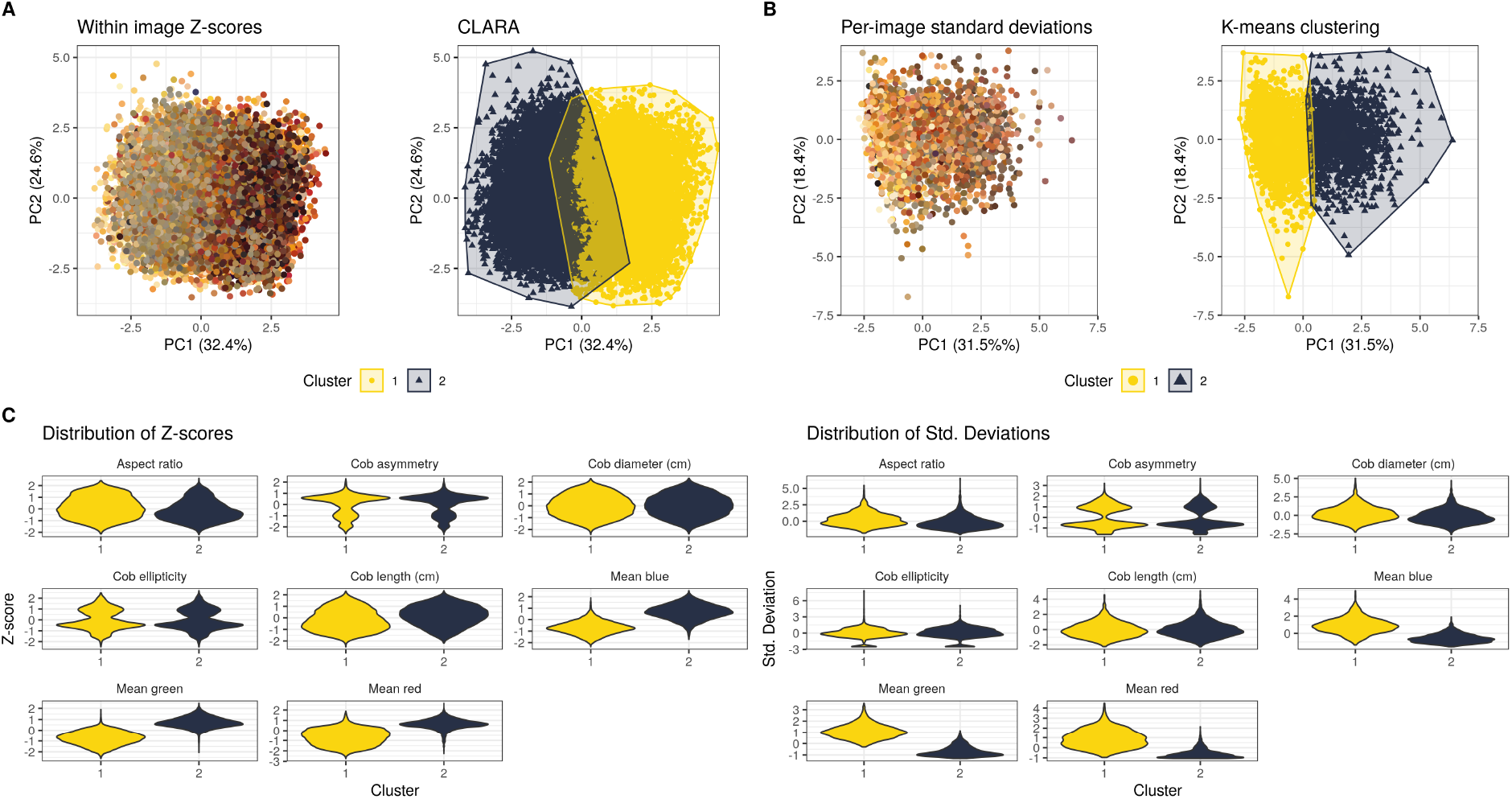
Clustering of individual images by their heterogeneity of maize cob traits within images. Clustering approaches with the extracted cob traits. (A) First two principal components showing the average color of individual cobs (*n* = 19, 867 cobs) (left) and average cob color per analyzed image (*n* = 3, 302 images) (right). The colors of each dot reflect the average RGB values (i.e., the color) of each cob, or image, respectively. (B) PCA plots showing clusters identified with CLARA (left) and *k* -means clustering (right). (C) Distribution of cob traits within each method and cluster.

## Discussion

Our comparison of three image segmentation methods showed Mask R-CNN to be superior to the classic image analysis method FelzSEG and WindowCNN for maize cob detection and segmentation. Given the recent success of Mask R-CNN for image segmentation in medicine or robotics, its application for plant phenotyping is highly promising as demonstrated in strawberry fruit detection for harvesting robots [38], orange fruit detection [39] and pomegranate tree detection [40]. Here we present another application of Mask R-CNN for maize cob instance segmentation and quantitative phenotyping in the context of genebank phenomics. In contrast to previous studies we performed a statistical analysis on the relative contribution of Mask R-CNN training parameters, and our application is based on more diverse and larger training image sets of 200 and 1,000 images. Finally, we propose a simple and rapid model updating scheme for applying the method on different maize cob image sets to make this method widely useful for cob phenotyping. The provided manuals offer a simple application and update of the deep learning model on custom maize cob datasets.

### Identifying optimal parameters for image segmentation

After optimizing various model parameters, the final Mask R-CNN model detected and segmented cobs and rulers very reliably with a very high *AP*@[.5: .95] score of 87.7, enabling accurate and fast extraction of cob features. Since such scores have not been reported for existing pipelines for maize cob annotation because they are mainly used for deep learning, we compared them to other contexts of image analysis and plant phenotyping where these parameters are available. Our score is higher than the original Mask R-CNN implementation on COCO with Cityscapes images [41], possibly due to a much smaller number of classes (2 versus 80) in our dataset. Depending on the backend network, the score of the original implementation ranged between 26.6 and 37.1. The maize cob score is also greater than 57.5 in the test set for pomegranate tree detection [40] and comparable to a score of 89.85 for strawberry fruit detection [38]. Although both maize cob and ruler detection and segmentation performed well, we observed minor inaccuracies in some masks. A larger training set did not improve precision and eliminate these inaccuracies, as the resolution of the mask branch in the Mask R-CNN framework may be too low, which could be improved by adding a convolutional layer of, for example, 56*×*56 pixel instead of the usual 28*×*28 pixel at the cost of longer computing time.

Mask R-CNN achieved higher correlation coefficients between true and predicted cob measurements than existing image analysis methods, which reported coefficients of *r* = 0.99 for cob length, *r* = 0.97 for cob diameter [14] and *r* = 0.93 for cob diameter [13]. Our Mask R-CNN achieved coefficients of *r* = 0.99 for cob diameter and *r* = 1 for cob length. Such correlations are a remarkable improvement considering that they were obtained with the highly diverse and inhomogeneous ImgOld and ImgNew image data (Table 8 and Supplementary Table S4), whereas previous studies used more homogeneous images with respect to color and shape of elite maize hybrid breeding material taken with uniform backgrounds. The high accuracy of Mask R-CNN indicate the advantage of the learning on specific cob and ruler patterns in deep learning.

Another feature of our automated pipeline is the simultaneous segmentation of cob and ruler, which allows pixel measurements to be instantly converted to centimeters and morphological measurements to be returned. Such an approach was also used by Makanza et al., [14], but no details on ruler measurements or accuracy of ruler detection were provided. The ability to detect rulers and cobs simultaneously is advantageous in a context where professional imaging equipment is not available, such as agricultural fields.

### Selection of training parameters to reduce annotation and training workload

Our Mask R-CNN workflow consists of annotating the data, training or updating the model, and running the pipeline to automatically extract features from the maize cobs. The most time-consuming and resource-intensive step was the manual annotation of cob images to provide labeled images for training, which took several minutes per image, but can be accelerated by supporting software [42]. In the model training step, model weights are automatically learned from the annotated images in an automated way, which is a major advantage over existing maize cob detection pipelines that require manual fine-tuning of parameters for different image datasets using operations such as thresholding, filtering, water-shedding, edge detection, corner detection, blurring and binarization [13, 14, 15].

Statistical analysis of each Mask R-CNN training parameters helps to reduce the amount of annotation and fine-tuning required (Tables 1 and 2). For example, there was no significant improvement on a large training set of 1,000 compared to 200 images, as learning on and segmenting of two object classes only seems to be a simple task for Mask R-CNN. Therefore, the significant amount of work involved in manual image annotation can be reduced if no more than 200 images need to be annotated. Since many training parameters did not have a strong impact on the final model result, this suggests that such parameters do not need to be fine-tuned. For example, using all layers instead of only the network heads (only the last part of the network involving the fully-connected layers) did not improve significantly the final detection result. Training image datasets with only a few object classes on network heads greatly reduces the runtime for model training.

### Technical equipment and computational resources for deep learning

The robustness of the Mask R-CNN approach imposes only simple requirements for creating images for both training and application purposes. RGB images taken with a standard camera are sufficient. In contrast, neural network training requires significant computational resources and is best performed on a high performance computing cluster or on GPUs with significant amounts of RAM. Training of the 90 different models (Table S6) was executed over 3 days, using 4 parallel GPUs on a dedicated GPU cluster. However, once the maize model is trained, model updating with only a few annotated images from new maize image data does not require a high performance computing infrastructure anymore, as in our case updating with 20 images was achieved in less than an hour on a normal workstation with 16 CPU threads and 64GB RAM.

Model updating with the pre-trained maize model on two different image datasets ImgCross and ImgDiv significantly improved the *AP*@[.5: .95] score for cob and ruler segmentation on the new images. The improvement was achieved despite additional features in the new image data that were absent from the training data. New features include rotated images, cobs in different orientation (horizontal instead of vertical) and different backgrounds (Figure 6). The advantage of a pre-trained maize model over the standard COCO model was independent of the image data set and achieved higher *AP*@[.5: .95] scores with a small number of epochs (Figure 5) because it saves training time for new image types, is widely applicable, and can be easily transferred to new applications for maize cob phenotyping. Importantly, the initial training set is not required for model updating. Our analyses indicate that only 10-20 annotated new images are required and the update can be limited to 50 epochs. The updated model can then be tested on the new image dataset, either by visual inspection of the detection or by annotating some validation images to obtain a rough estimate of the *AP*@[.5: .95] score. The phenotypic traits can then be extracted by the included post-processing workflow, which itself only needs to be modified if additional parameters are to be implemented.

The runtime of the pipeline after model training is very fast. Image segmentation with the trained Mask R-CNN model and parameter estimation of eight cob traits took on average of 3.6 seconds per image containing an average of six cobs. This time is shorter than previously published pipelines (e.g., 13 seconds per image in [13]), although it should be noted that any such comparisons are not based on the same hardware and the same set of traits. For example, the pipeline for three dimensional cob phenotyping performs a flat projection of the surface of the entire cob, but is additionally capable of annotating individual cob kernels and the total time for analyzing a single cob is 5-10 minutes [15]. The ear digital imaging (EDI) pipeline of Makanza et al. [14] processes more than 30 unthreshed ears at the same time and requires more time per image at 10 seconds, but also extracts more traits. However, this pipeline was developed on uniform and standardized images and does not involve a deep learning approach to make it generally applicable.

### Application of the Mask R-CNN pipeline for genebank phenomics

To demonstrate the utility of our pipeline, we applied it to original images of maize cobs from farmer’s fields during the establishment of the official maize genebank in Peru in the 1960s and 1970s (ImgOld) and to more recent photographs taken during the regeneration of existing maize material in 2015 (ImgNew). The native maize diversity of Peru was divided into individual landraces based mainly on cob traits. Our interest was to identify genebank accessions with high or low diversity of cob traits within accessions to classify accessions as ‘pure’ representatives of a landrace or as accessions with high levels of native genetic diversity, evidence of recent gene flow, or random admixture of different landraces. We used two different approaches to characterize the amount of variation for each trait within the accessions based on the eight traits measured by our pipeline. Unsupervised clustering of variance measure identified two groups of accessions that differed in their overall level of variation. The distribution of normalized variance parameters (Z-scores and standard deviations) within both groups indicate that variation in cob color has the strongest effect on variation within genebank accessions, suggesting that cob color is more variable that morphometric characters like cob length or cob diameter. This information is useful for subsequent studies, in terms of the relationship between genetic and phenotypic variation in native maize diversity, the geographic patterns of phenotypic variation within landraces, or the effect of seed regeneration during *ex situ* conservation on phenotypic diversity, which we are currently investigating in a separate study.

## Conclusion

We present the successful application of deep learning by Mask R-CNN to maize cob segmentation in the context of genebank phenomics by developing a pipeline written in Python for a large-scale image analysis of highly diverse maize cobs. We also developed a post-processing workflow to automatically extract measurements of eight phenotypic cob traits from cob and ruler masks obtained with Mask R-CNN. In this way, cob parameters were extracted from 19,867 individual cobs with a fast automated pipeline suitable for high-throughput phenotyping. Although the Mask R-CNN model was developed based on native maize diversity of Peru, the model can be easily used and updated for additional image types in contexts like the genetic mapping of cob traits or in breeding programs. It therefore is of general applicability in maize breeding and research and for this purpose, we provide simple manuals for maize cob detection, parameter extraction and deep learning model updating. Future developments of the pipeline may include linking it to mobile phenotyping devices for realtime measurements in the field and using the large number of segmented images to develop refined models for deep learning, for example, to estimate additional parameters such a row numbers or characteristics of individual cob kernels.

## Materials and Methods

### Plant material

The plant material used in this study is based on 2,484 genebank accessions of 24 Peruvian maize landraces collected from farmer’s fields in the 1960s and 1970s, which are stored the Peruvian maize genebank hosted at the Universidad Agraria La Molina (UNALM), Peru. These accessions originate from the three different ecogeographical environments (coast, highland and rainforest) present in Peru and therefore represent a broad sample of Peruvian maize diversity.

### Image data of maize cobs

All accessions were photographed during their genebank registration. An image was taken with a set of 1-12 maize cobs per accession laid out side by side with a ruler and accession information. Because the accessions were collected over several years, the images were not taken under the same standardized conditions of background, rulers and image quality. Prints of these photographs were stored in light-protected cupboards of the genebank and were digitized with a flatbed scanner in 2015 and stored as PNG files without further image processing. In addition, all genebank accession were regenerated in 2015 at three different locations reflecting their ecogeographic origin and the cobs were photographed again with modern digital equipment under standardized conditions and also stored as PNG images. The image data consist thus consist of 1,830 original (ImgOld) and 1,619 new (ImgNew) images for a total of 3,449 images. Overall, the images show a high level of variation due to technical and genetic reasons, which are outlined in Figure 8. These datasets were used for training and evaluation of the image segmentation methods.

**Figure 8:**
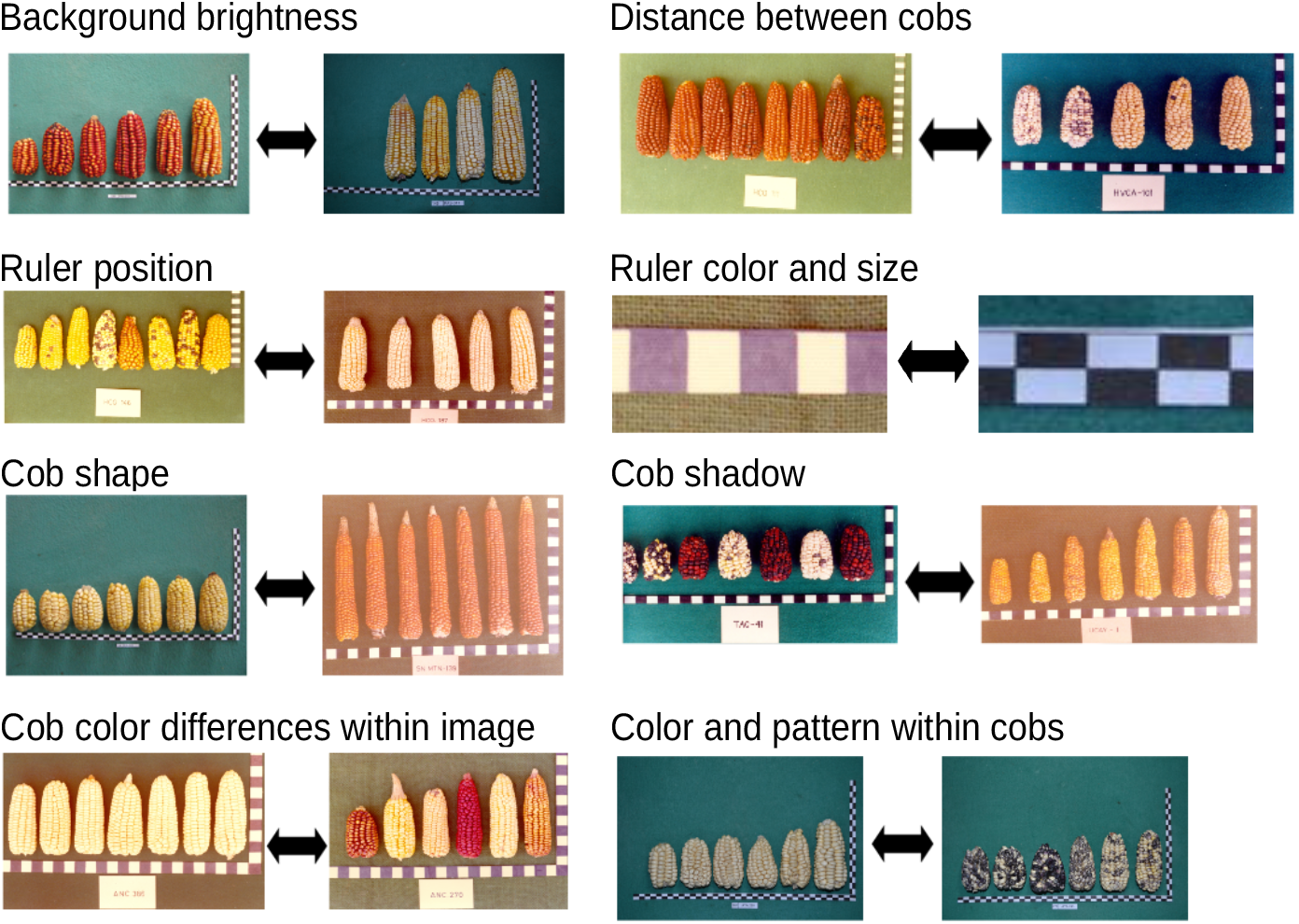
Variability of image properties among the complete dataset (containing ImgOld and ImgNew)

Passport information available for each accession and their assignment to the different landraces is provided in Table S5. All images were re-scaled to a size of 1000×666 pixels with OpenCV, version 3.4.2; [43].

We used two different datasets for updating the image segmentation models and evaluating their robustness. The ImgCross image dataset contains images of maize cobs and spindles derived from a cross of Peruvian landraces with a synthetic population generated from European elite breeding material and therefore reflects genetic segregation in the F2 generation. The images were taken with digital camera at the University of Hohenheim under standardized conditions and differ from the other data sets by a uniform green background, a higher resolution 3888×2592 pixels (no re-sizing), a variable orientation of the cobs, orange labels and differently colored squares instead of a ruler.

A fourth set of images (ImgDiv) was obtained mainly from publicly available South American maize genebank catalogs and from special collections available as downloadable figures on the internet. The ImgDiv data vary widely in terms of number and color of maize cobs, image dimensions and resolution, number, position and orientation of cobs. Some images also contain rulers as in ImgOld and ImgNew.

### Software and methods for image analysis

Image analysis was mainly performed on a workstation running Ubuntu 18.04 LTS and the analysis code was written in Python (version 3.7; [44]) for all image operations. OpenCV (version 3.4.2; [43]) was used to perform basic image operations like resizing and contour finding.

For WindowCNN and Mask R-CNN, deep learning was performed with the Tensorflow (version 1.5.0; [45]) and Keras (version 2.2.4; [46]) libraries. In Mask R-CNN, the framework [47] from the matterport implementation (https://github.com/matterport/Mask_RCNN) was used and adapted to the requirements of the maize cob image datasets. Statistical analyses for evaluating the contribution of different parameters in Mask R-CNN and for the clustering of the obtained cob traits was carried out with R version 3.6.3 [48].

We tested three different approaches (FelzSEG, WindowCNN and Mask R-CNN) for cob and ruler detection and image segmentation. Details on their implementation and comparison can be found in the Supplementary Text, but our approach is briefly described below.

For image analysis using traditional approaches, we first applied various tools such as filtering, water-shedding, edge detection and corner detection to representative subsets of ImgOld and ImgNew. The best segmentation results were obtained with the graph-based Felzenszwalb-Huttenlocher image segmentation algorithm [49] implemented in the Python scikit-image library version 0.16.2 [50] and the best ruler detection with the naive Bayes Classifier, implemented in the PlantCV library [51]. The parameters had to be manually fine-tuned for each of the two image datasets.

Too evaluate deep learning, we used a windows-based (WindowCNN) and a Mask R convolutional neural network (Mask R-CNN), both of which require training on annotated and labeled image data. Convolutional Neural Networks [52] (CNN) are known to be the most powerful feature extractors and their popularity for image classification dates back to the ImageNet classification challenge, which was won by the architecture AlexNet [53]. Generally, a CNN consists of 3 different layer types, which are subsequently connected: Convolutional layers, Pooling Layers and Fully-Connected (FC) Layers. In a CNN for cob detection classes ‘cob’ and ‘ruler’ can be learned as a feature using deep learning, which provides maize cob feature extraction independent of the challenges in diverse images like scale, cob color, cob shape, background color and contrast.

Since our goal was to localize and segment the cobs within the image, we first used sliding window CNN (WindowCNN), which passes parts of an image to a CNN at a time and returns the probability that it contains a particular object class. Sliding windows have been used in plant phenotyping to detect plant segments [54, 55]. Our implementation of WindowCNN is described in detail in Chapter S**??**.

Since sliding window CNN have low accuracy and very long runtime, feature maps are used to filter out putative regions of interest on which boxes are refined around objects. Mask R-CNN [47] is the most recent addition to the family of R-CNN [56] and includes a Region Proposal Network (RPN) to reduce the number of bounding boxes by passing only *N* region proposals that are likely to contain some object to a detection network block. The detection network generates the final object localizations along with the appropriate classes from the RPN proposals and the appropriate features from the feature CNN. Mask R-CNN extends a Fast R-CNN [57] with a mask branch of two additional convolutional layers that perform additional instance segmentation and return a pixel-wise mask for each detected object containing a bounding box, a segmentation mask and a class label.

### Implementation of Mask R-CNN to detect maize cobs and rulers

The training image data (200 or 1,000 images) were randomly selected from the two datasets ImgOld and ImgNew to achieve maximum diversity in terms of image properties (Table 8 and Supplementary Table S4). Both subsets were each randomly divided into a training set (75%) and a validation set (25%). Both image subsets were annotated using VGG Image Annotator (via; version 2.0.8 [58]). A pixel-precise mask was drawn by hand around each maize cob (Supplementary Figure S2). The ruler was labeled with two masks, one for the horizontal part and one for the vertical part, which facilitates later prediction of the bounding boxes of the ruler compared to annotating the entire ruler element as one mask. Each mask was labeled as “cob” or “ruler”, and the annotations for training and validation sets were exported separately as JSON files.

The third step consisted of model training on multiple GPUs using a standard tensorflow implementation of Mask R-CNN for maize cob and ruler detection. We used the pre-trained weights of the COCO model, which is the standard model [47] derived from training on the MS COCO dataset [37], in the layout of resnet 101 (transfer learning). The original Mask R-CNN implementation was modified by adding two classes for cob and ruler in addition to the background class. Instead of saving all models after each training epoch, only the best model with the least validation loss was saved to save memory. For training the Mask R-CNN models, we used Tesla K80 GPUs with 12 GB RAM each on the BinAC GPU cluster at the University of Tü bingen [59].

We trained 90 different models with different parameter settings (Supplementary Table S6) on both image datasets. The learning rate parameter *learningrate* was set to vary from 10^*−*3^, as in the standard implementation, to 10^*−*5^, since models with smaller datasets often suffer from overfitting, which may require smaller steps in learning the model parameters. Training was performed over 15, 50, or 200 epochs (*epochsoverall*) to capture potential overfitting issues. The parameter *epochs*.*m* distinguishes between training only the heads, or training the heads first, followed by training on the complete layers of resnet101. The latter requires more computation time, but offers the possibility to fine tune not only the heads, but all the layers to obtain a more accurate detection. Also, the mask loss weight (*masklossweight*) was given the value of 1, as in the default implementation, or 10, which means a higher focus on reducing mask loss. Also, the monitor metric (*monitor*) for the best model checkpoint was set to vary between the default validation loss and the mask validation loss. The latter option was tested to optimize preferentially for mask creation, which is usually more challenging than determining object class, bounding box loss, etc. The use of the minimask (*minimask*) affects the accuracy of mask creation and in the default implementation consists of a resizing step before the masks are forwarded by the CNN during the training process.

The performance of these models for cob and ruler detection was evaluated by the IoU (Intersection over Union) score or Jaccard index [60], which is the most popular metric to evaluate the performance of object detectors. The IoU score between a predicted and a true bounding box is calculated by

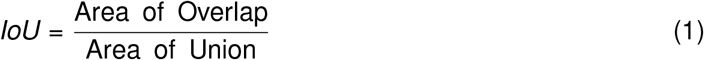

The most common threshold for IoU is 50% or 0.5. With IoU values above 0.5, the predicted object is considered as true positive (TP), else as a false positive (FP). Precision is calculated by

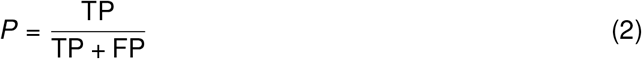

The average precision (AP) was calculated by averaging *P* over all ground-truth objects of all classes in comparison to their predicted boxes, as demonstrated in various challenges and improved network architectures [61, 62, 63].

Following the primary challenge metric of the COCO dataset [64], the goodness of our trained models was also scored by *AP*@[.5: .95], sometimes also just called AP, which is the average AP over different IoU thresholds from 50% to 95% in 5% steps. In contrast to usual object detection models where IoU/AP metrics are calculated for boxes, in the following IoU relates to the masks [41], because this explores the performance of instance segmentation. We performed an ANOVA with 90 model results scores to evaluate the individual impact of the parameters on the *AP*@[.5: .95] score. Logit transformation was applied to fit the assumptions of heterogeneity of variance and normal distribution (Supplementary Figure S3). Model selection was carried out including parameters *learningrate* (10^*−*3^, 10^*−*4^, 10^*−*5^, *epochs*.*m* (1:only heads, 2:20 epochs heads, 3:10 epochs heads; for the rest all model layers trained), *epochsoverall* (15, 50, 200), *masklossweight* (1,10), *monitor* (val loss, mask val loss) and *minimask* (yes, no). Also all two-way interactions were included in the model, dropping non-significant interactions first and then non-significant main effects if none of their interactions were significant.

These results allow to formulate the following final model to describe contributions of the parameters on Mask R-CNN performance:

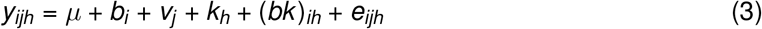

where *µ* is the general effect, *b*_*i*_ the effect of the *i* -th minimask, *v*_*j*_ the effect of the *j* -th overall number of epochs, *k*_*h*_ the effect of the *h*-th training set size, (*bk*)_*ih*_ the interaction effect between the number of epochs and the training set size and *e*_*ijh*_ the random deviation associated with *y*_*ijh*_. We calculated

ANOVA tables, back-transformed lsmeans and contrasts (confidence level of 0.95) for the significant influencing variables. As last step of model training, we set up a workflow with the best model as judged by its *AP*@[.5: .95] score and performed random checks whether objects were detected correctly.

### Workflow for model updating with new pictures

To investigate the updating ability of Mask R-CNN on different maize cob image datasets, we annotated additionally 150 images (50 training, 100 validation images) from each of the ImgCross and ImgDiv datasets. For ImgCross, the high resolution of 3888 *×* 2592 pixels was maintained, but 75% of the images were rotated (25% by 90^*°*^, 25% by 180^*°*^, and 25% by 270^*°*^) to increase diversity. The corn cob spindles on these images were also labeled as cobs and the colored squares were labeled as rulers. The ImgDiv images were left at their original resolution and annotated with the cob and ruler classes.

The model weights of the best model (M104) obtained by training with ImgOld and ImgNew were used as initial weights and updated with ImgCross and ImgDiv images. Based on the statistical analysis, optimal parameter levels of the main parameters were used and only the network heads were trained with a learning rate of 10^*−*4^ for 50 epochs without the minimum mask. Training was performed with different randomly selected sets (10, 20, 30, 40, and 50 images) to evaluate the influence of the number of images on the quality of model updating. For each training run, all models with an improvement step in validation loss were saved, and the *AP*@[.5:.95] score was calculated for each of them. For comparison, all combinations of models were also trained with the standard COCO weights.

### Statistical analysis of model updating results

To evaluate the influence of the data set, the starting model, and the size of the training set, an ANOVA was performed on the data set of *AP*@[.5: .95] from all epochs and combinations. Logit transformation was applied to meet the assumptions of heterogeneity of variance and normal distribution. Epoch was included as a covariate. Forward model selection was performed using the parameters *dataset* (ImgCross, ImgDiv), *starting model* (COCO, pre-trained maize model), and *training set size* (10, 20, 30, 40, 50). All two-way and three-way parameter interactions were included in the model. Because the three-way interaction was not significant, the significant two-way interactions and significant main effects were retained in the final model, which can be denoted as follows:

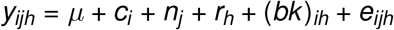

where

*µ* = general effect

*c*_*i*_ = effect of the i-th dataset

*n*_*j*_ = effect of the j-th starting model

*k*_*h*_ = effect of the h-th training set size

(*cn*)_*ih*_ = interaction effect between the dataset and the starting model

(*nk*)_*jh*_ = interaction effect between the starting model and the training set size

*e*_*ijh*_ = random deviation associated with *y*_*ijh*_

ANOVA tables, back-transformed lsmeans (Supplementary Tables S7 and S8) and contrasts (confidence level of 0.95) for the significant influencing variables were calculated.

### Post-processing of segmented images for automated measurements and phenotypic trait extraction

Mask R-CNN images are post-processed with an automated pipeline to extract phenotypic traits of interest such as cob shape or cob color descriptors (Figure 9). The Mask R-CNN model returns a list of labeled masks, which are separated into cob and ruler masks for subsequent analysis. Contour detection is applied to binarized ruler masks to identify individual black or white ruler elements, whose length in pixel is then average for elements of a ruler to obtain a pixel value per cm for each image. Length and diameter of cob masks are then converted from pixel into cm values using the average ruler lengths. The cob masks are also used to calculate the mean RGB color of each cob. In contrast to a similar approach by Miller et al. [13], who sampled pixels from the middle third of cobs for RGB color extraction, we used the complete cob mask because kernel color was variable throughout the cob in highly diverse image data. We also used the complete cob mask to extract cob shape parameters that include asymmetry and ellipticity similar to a previous study of avian eggs [65], who characterized egg shape diversity using the morphometric equations of Baker [66]. Since our image data contained a high diversity of maize cob shapes we reasoned that shape parameters like asymmetry and ellipticity are useful for a morphometric description of maize cob diversity. Overall the following phenotypic traits were extracted from almost 19,867 cobs: Diameter, length, aspect ratio (length/diameter), asymmetry, ellipticity and mean RGB color separated by red, green, blue channels. Our pipeline returned all cob masks for later analysis of additional parameters as .jpg images.

**Figure 9:**
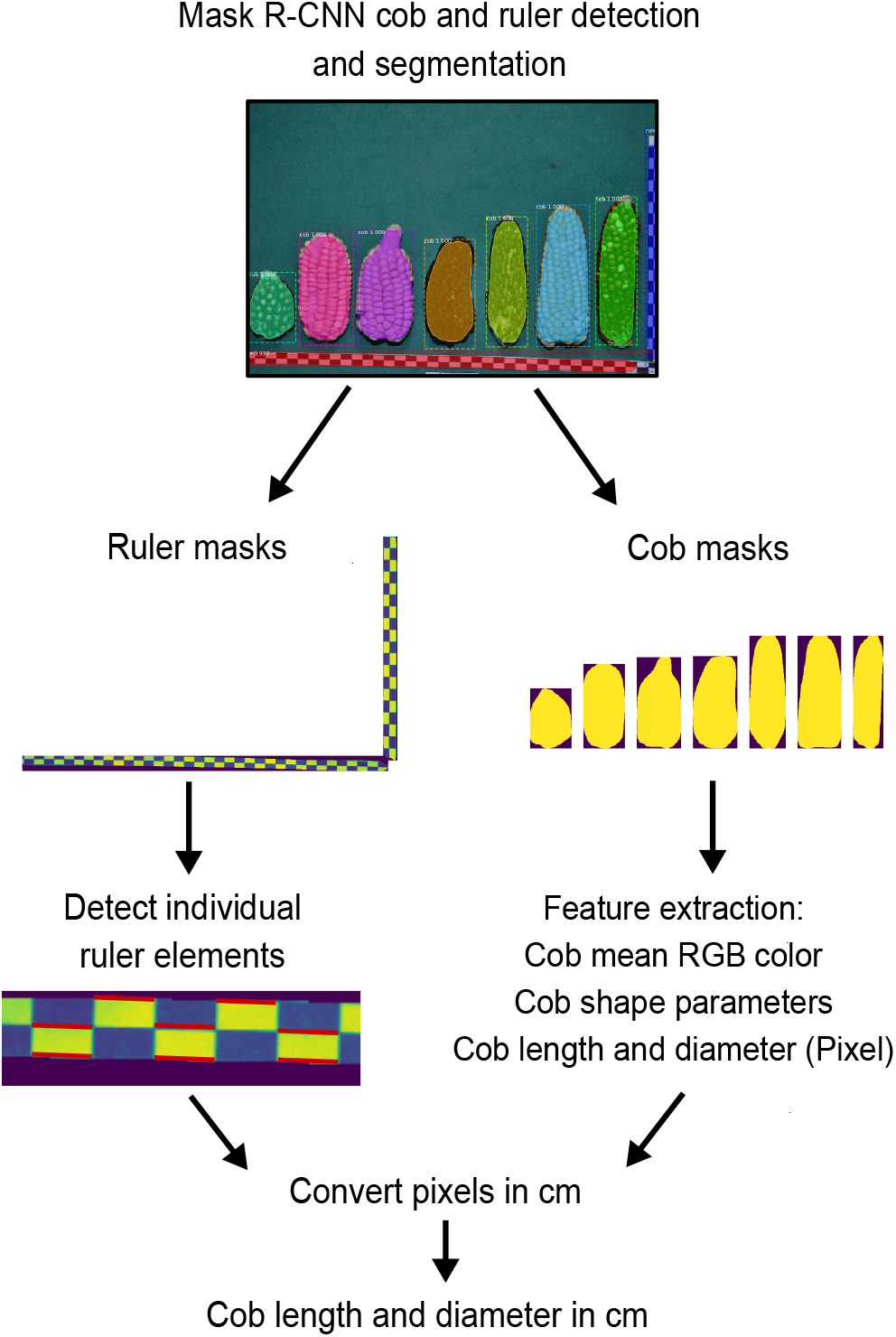
Post-processing of segmented images using a Mask R-CNN workflow that analyses segments labeled as ‘cob’ and ‘ruler’ to extract the parameters cob length, diameter, mean RGB color,and shape parameters ellipticity and asymmetry. Cob length and diameter measures in pixels are converted to cm values by measuring the contours of single ruler elements.

### Quantitative comparison between FelzSEG, WindowCNN and Mask R-CNN

For quantitative comparisons between the three image segmentation methods, a subset of 50 images from ImgOld and 50 images from ImgNew were randomly selected. None of the images were included in the training data from WindowCNN or Mask R-CNN, and the subset is unbiased against the training data. True measurements of cob length and diameter were obtained using the annotation tool *via* [58]. Individual cob dimensions per image could not be directly compared to predicted cob dimensions because FelzSEG and WindowCNN often contained multiple cobs in a box or certain cobs were contained in multiple boxes. Therefore, the mean of the predicted cob width and length per image was calculated for each approach, penalizing incorrectly predicted boxes. Pearson correlation was calculated between the true and predicted mean diameter and length of the cob per image separately for the ImgOld and ImgNew sets.

### Unsupervised clustering to detect images with high cob diversity

To identify genebank accessions with high phenotypic diversity in ImgOld and ImgNew images, we used two different unsupervised clustering methods. In the first approach, individual cob features (width, length, asymmetry, ellipticity, and mean RGB values) were scaled after their extraction from the images. The Z-score of each cob was calculated as 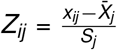, where *Z*_*ij*_ is the Z-score of the *i* th cob in the *j* th image, *x*_*ij*_ is a measurement of the *i* th cam of the *j* th image, and 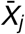 and *S*_*j*_ are the mean and are the standard deviation of the *j* -th image, respectively. The scaled dataset was analyzed using CLARA (Clustering LARge Applications) as described in the *cluster* R package [67]. The optimal cluster number was determined by the average silhouette method implemented in the R package factoextra [68].

In the second approach, we used the standard deviations of individual measurements within each each image (*S*_*j*_) as input for clustering. The standard deviations of each image were centered and standardized so that the values obtained for all images were on the same scale. This dataset was then clustered with *k* -means and the number of clusters, *k*, was determined using the gap statistic [69], which compares the sum of squares within clusters to the expectation under a zero reference distribution.

## Abbreviations

*AP*@[.5: .95]: AP@[IoU=0.50:0.95] sometimes also called mAP.
CLARA: Clustering Large Applications
RPN: Region Proposal Network

## Supplementary Information

- Supplementary Tables and Figures
- Supplementary Text

## Declarations

## Acknowledgments

We are grateful to Gilberto Garcia for scanning and photographing the maize genebank accessions at UNALM, Emilia Koch for annotating the images, and Hans-Peter Piepho for statistical advice.

## Author contributions

LK and KS designed the study. LK performed the image analysis, implemented FelzSEG, WindowCNN and Mask R-CNN on the datasets, developed the model updating and carried out the statistical analyses. MCA conducted the multivariate analysis of phenotypic cob data. RB coordinated and designed the acquisition of the maize photographs. LK and KS wrote the manuscript. All authors read, revised and agreed on the manuscript.

## Availability of data and materials

- Image files and annotations: http://doi.org/10.5281/zenodo.4587304
- Deep learning model and manuals with codes for custom detections and model updating: https://gitlab.com/kjschmidlab/deepcob

### Ethics approval and consent to participate

Not applicable.

### Consent for publication

Not applicable.

