## Supplementary Figures and Tables for "DeepCob: Precise and high-throughput analysis of maize cob geometry using deep learning with an application in genebank phenomics"

#### Supplementary tables

Supplementary Table S1: Training results of the 90 *Mask R-CNN* models. AP@[.5 : .95] scores of the 90 models with different training parameters, for training sets of 200 and 1000 images.

| Model | 200 images | 1000 images | Model | 200 images | 1000 images |
|---|---|---|---|---|---|
| M01 | 54.60 | 57.52 | M101 | 86.56 | 79.67 |
| M02 | 57.74 | 38.06 | M102 | 84.68 | 81.14 |
| M03 | 67.95 | 46.42 | M103 | 79.72 | 76.17 |
| M04 | 65.64 | 44.12 | M104 | 86.74 | 80.02 |
| M05 | 63.77 | 52.10 | M105 | 81.92 | 79.14 |
| M06 | 29.96 | 50.78 | M106 | 79.39 | 78.81 |
| M07 | 55.21 | 52.76 | M107 | 86.56 | 69.68 |
| M08 | 59.18 | 66.05 | M108 | 83.89 | 80.05 |
| M09 | 69.08 | 21.60 | M109 | 79.95 | 76.98 |
| M11 | 46.38 | 45.70 | M111 | 81.18 | 78.88 |
| M12 | 73.69 | 55.21 | M112 | 73.76 | 77.21 |
| M13 | 40.08 | 46.51 | M113 | 74.94 | 75.19 |
| M14 | 25.75 | 44.06 | M114 | 84.80 | 83.78 |
| M15 | 41.84 | 46.67 | M115 | 84.80 | 78.02 |
| M16 | 48.73 | 59.71 | M116 | 81.91 | 77.96 |
| M17 | 68.58 | 34.37 | M117 | 84.24 | 77.25 |
| M18 | 51.54 | 56.15 | M118 | 83.43 | 76.56 |
| M19 | 59.78 | 61.71 | M119 | 82.80 | 81.56 |
| M21 | 61.83 | 54.36 | M121 | 80.75 | 80.28 |
| M22 | 58.53 | 65.13 | M122 | 78.59 | 75.54 |
| M23 | 48.76 | 10.49 | M123 | 68.42 | 73.25 |
| M24 | 69.17 | 38.14 | M124 | 86.56 | 79.10 |
| M25 | 62.00 | 21.68 | M125 | 84.83 | 78.60 |
| M26 | 63.47 | 29.07 | M126 | 81.89 | 79.21 |
| M27 | 41.71 | 38.36 | M127 | 82.00 | 79.74 |
| M28 | 75.96 | 43.72 | M128 | 85.87 | 79.04 |
| M29 | 59.29 | 64.24 | M129 | 62.80 | 79.84 |
| M31 | 28.11 | 54.18 | M01 | 83.09 | 81.76 |
| M32 | 63.71 | 37.46 | M01 | 82.36 | 83.13 |
| M33 | 51.76 | 52.95 | M01 | 73.26 | 73.81 |
| M34 | 68.33 | 43.32 | M01 | 86.13 | 78.73 |
| M35 | 72.48 | 50.09 | M01 | 84.78 | 79.35 |
| M36 | 50.48 | 66.79 | M01 | 81.95 | 81.86 |
| M37 | 58.66 | 35.87 | M01 | 85.23 | 79.60 |
| M38 | 47.44 | 54.04 | M01 | 84.87 | 78.91 |
| M39 | 44.80 | 60.39 | M01 | 82.71 | 83.42 |
| M41 | 5.57 | 57.13 | M131 | 77.30 | 76.84 |
| M42 | 52.13 | 54.41 | M132 | 76.41 | 57.45 |
| M43 | 35.00 | 56.35 | M133 | 53.91 | 70.78 |
| M44 | 52.04 | 30.45 | M134 | 84.86 | 84.31 |
| M45 | 42.20 | 43.72 | M135 | 83.74 | 75.36 |
| M46 | NA | 34.52 | M136 | 74.29 | 75.66 |
| M47 | 43.45 | 56.20 | M137 | 85.03 | 78.15 |
| M48 | 30.75 | 32.01 | M138 | 76.27 | 83.45 |
| M49 | 31.50 | 62.78 | M139 | 69.52 | 70.62 |

Supplementary Table S2: P-values of lsmeans for Mask R-CNN model parameter *minimask* with parameter values *yes* or *no*.

| Contrast | Estimate | SE | df | t ratio | p-value |
|---|---|---|---|---|---|
| no - yes | 1.4 | 0.07 | 170 | 19.61 | $<$0.0001 |

Supplementary Table S3: P-values of lsmeans for Mask R-CNN model parameters *trainingdata* $\times$ *epochsoverall*. Parameter values for training data are 200 or 1,000 images, and for epochsoverall 15, 20, 200.

| Contrast | Estimate | SE | df | t ratio | p-value |
|---|---|---|---|---|---|
| 200 images; 15 epochs vs. 200 images; 50 epochs | 0.36 | 0.13 | 170 | -4.39 | 0.0003 |
| 200 images; 15 epochs vs. 200 images; 200 epochs | 0.33 | 0.16 | 170 | -4.42 | 0.0003 |
| 1000 images; 50 epochs vs. 200 images; 200 epochs | 0.41 | 0.13 | 170 | -2.96 | 0.0407 |
| 1000 images; 15 epochs vs. 200 images; 200 epochs | 0.38 | 0.16 | 170 | -2.95 | 0.0413 |
| 1000 images; 50 epochs vs. 200 images; 50 epochs | 0.44 | 0.09 | 170 | -2.77 | 0.0682 |
| 1000 images; 15 epochs vs. 200 images; 50 epochs | 0.42 | 0.13 | 170 | -2.61 | 0.0995 |
| 200 images; 15 epochs vs. 1000 images; 50 epochs | 0.42 | 0.13 | 170 | -2.48 | 0.1359 |
| 1000 images; 200 epochs vs. 200 images; 200 epochs | 0.4 | 0.16 | 170 | -2.47 | 0.1391 |
| 200 images; 50 epochs vs. 1000 images; 200 epochs | 0.57 | 0.13 | 170 | 2.02 | 0.3334 |
| 200 images; 15 epochs vs. 1000 images; 200 epochs | 0.42 | 0.16 | 170 | -1.98 | 0.3553 |
| 1000 images; 15 epochs vs. 200 images; 15 epochs | 0.56 | 0.16 | 170 | 1.51 | 0.6584 |
| 200 images; 50 epochs vs. 200 images; 200 epochs | 0.47 | 0.13 | 170 | -1 | 0.9169 |
| 1000 images; 15 epochs vs. 1000 images; 50 epochs | 0.48 | 0.13 | 170 | -0.66 | 0.9861 |
| 1000 images; 15 epochs vs. 1000 images; 200 epochs | 0.48 | 0.16 | 170 | -0.48 | 0.9967 |
| 1000 images; 50 epochs vs. 1000 images; 200 epochs | 0.5 | 0.13 | 170 | 0.07 | 1 |

Supplementary Table S4: Variability of image properties specifically between ImgOld and ImgNew, which were caused only by different imaging conditions. The images taken in ImgNew were generally more standardized than in ImgOld.

| Source of variation | ImgOld | ImgNew | Examples |
|---|---|---|---|
| Image resolution | variable resolution | standardized resolution | |
| Background color | diverse | dark blue only | |
| Incomplete cobs | present | absent | 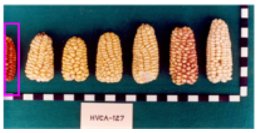 |
| Ruler scale horizontal<br>Ruler scale vertical<br>Ruler type | 2 cm<br>2 cm<br>unstacked | 1 cm<br>0.5 cm<br>stacked | 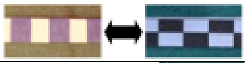 |
| Figure orientation | landscape | landscape or portrait | 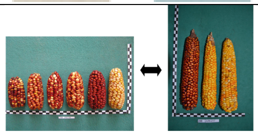 |

Supplementary Table S5: Number of populations and respective environment represented per department in ImgOld and ImgNew.

| | | Number of accessions | |
|---|---|---|---|
| Maize landrace | Environment | ImgOld | ImgNew |
| Rainforest | Amazonas | 23 | 10 |
| | Huanuco | 98 | 28 |
| | Loreto | 29 | 31 |
| | M.Dios | 31 | 33 |
| | San Martin | 89 | 105 |
| | Ucayali | 29 | 20 |
| Highland | Ancash | 316 | 259 |
| | Apurimac | 175 | 158 |
| | Ayacucho | 79 | 128 |
| | Cajamarca | 75 | 34 |
| | Cusco | 111 | 80 |
| | Huancavelica | 73 | 78 |
| | Junin | 85 | 160 |
| | Pasco | 25 | 20 |
| | Puno | 14 | 12 |
| Coast | Arequipa | 80 | 68 |
| | Ica | 50 | 27 |
| | Lambayeque | 120 | 99 |
| | Libertad | 113 | 93 |
| | Lima | 58 | 54 |
| | Mgua | 8 | 5 |
| | Piura | 91 | 88 |
| | Tacna | 39 | 19 |
| | Tumbes | 16 | 10 |
| | Total | 1827 | 1619 |

Supplementary Table S6: Parameter combinations of the 90 trained *Mask R-CNN* models. 90 trained *Mask R-CNN* models differing in the parameters *learningrate*, *epochs.m*, *epochsoverall*, *masklossweight* and *monitor*. Each model was repeated with and without the use of a mini mask (*minimask*).

| model | *learningrate* | *epochsoverall* | *epochs.m* | *masklossweight* | *monitor* | *minimask* |
|---|---|---|---|---|---|---|
| M01 | 1e-3 | 200 | 1 | 10 | val loss | yes |
| M02 | 1e-4 | 200 | 1 | 10 | val loss | yes |
| M03 | 1e-5 | 200 | 1 | 10 | val loss | yes |
| M04 | 1e-3 | 200 | 3 | 10 | val loss | yes |
| M05 | 1e-4 | 200 | 3 | 10 | val loss | yes |
| M06 | 1e-5 | 200 | 3 | 10 | val loss | yes |
| M07 | 1e-3 | 200 | 2 | 10 | val loss | yes |
| M08 | 1e-4 | 200 | 2 | 10 | val loss | yes |
| M09 | 1e-5 | 200 | 2 | 10 | val loss | yes |
| M11 | 1e-3 | 50 | 1 | 1 | mask val loss | yes |
| M12 | 1e-4 | 50 | 1 | 1 | mask val loss | yes |
| M13 | 1e-5 | 50 | 1 | 1 | mask val loss | yes |
| M14 | 1e-3 | 50 | 3 | 1 | mask val loss | yes |
| M15 | 1e-4 | 50 | 3 | 1 | mask val loss | yes |
| M16 | 1e-5 | 50 | 3 | 1 | mask val loss | yes |
| M17 | 1e-3 | 50 | 2 | 1 | mask val loss | yes |
| M18 | 1e-4 | 50 | 2 | 1 | mask val loss | yes |
| M19 | 1e-5 | 50 | 2 | 1 | mask val loss | yes |
| M21 | 1e-3 | 50 | 1 | 1 | val loss | yes |
| M22 | 1e-4 | 50 | 1 | 1 | val loss | yes |
| M23 | 1e-5 | 50 | 1 | 1 | val loss | yes |
| M24 | 1e-3 | 50 | 3 | 1 | val loss | yes |
| M25 | 1e-4 | 50 | 3 | 1 | val loss | yes |
| M26 | 1e-5 | 50 | 3 | 1 | val loss | yes |
| M27 | 1e-3 | 50 | 2 | 1 | val loss | yes |
| M28 | 1e-4 | 50 | 2 | 1 | val loss | yes |
| M29 | 1e-5 | 50 | 2 | 1 | val loss | yes |
| M31 | 1e-3 | 50 | 1 | 10 | val loss | yes |
| M32 | 1e-4 | 50 | 1 | 10 | val loss | yes |
| M33 | 1e-5 | 50 | 1 | 10 | val loss | yes |
| M34 | 1e-3 | 50 | 3 | 10 | val loss | yes |
| M35 | 1e-4 | 50 | 3 | 10 | val loss | yes |
| M36 | 1e-5 | 50 | 3 | 10 | val loss | yes |
| M37 | 1e-3 | 50 | 2 | 10 | val loss | yes |
| M38 | 1e-4 | 50 | 2 | 10 | val loss | yes |
| M39 | 1e-5 | 50 | 2 | 10 | val loss | yes |
| M41 | 1e-3 | 15 | 1 | 10 | val loss | yes |
| M42 | 1e-4 | 15 | 1 | 10 | val loss | yes |
| M43 | 1e-5 | 15 | 1 | 10 | val loss | yes |
| M44 | 1e-3 | 15 | 3 | 10 | val loss | yes |
| M45 | 1e-4 | 15 | 3 | 10 | val loss | yes |
| M46 | 1e-5 | 15 | 3 | 10 | val loss | yes |
| M47 | 1e-3 | 15 | 2 | 10 | val loss | yes |
| M48 | 1e-4 | 15 | 2 | 10 | val loss | yes |
| M49 | 1e-5 | 15 | 2 | 10 | val loss | yes |
| M101 | 1e-3 | 200 | 1 | 10 | val loss | no |
| M102 | 1e-4 | 200 | 1 | 10 | val loss | no |
| M103 | 1e-5 | 200 | 1 | 10 | val loss | no |
| M104 | 1e-3 | 200 | 3 | 10 | val loss | no |
| M105 | 1e-4 | 200 | 3 | 10 | val loss | no |
| M106 | 1e-5 | 200 | 3 | 10 | val loss | no |
| M107 | 1e-3 | 200 | 2 | 10 | val loss | no |
| M108 | 1e-4 | 200 | 2 | 10 | val loss | no |
| M109 | 1e-5 | 200 | 2 | 10 | val loss | no |
| M111 | 1e-3 | 50 | 1 | 1 | mask val loss | no |
| M112 | 1e-4 | 50 | 1 | 1 | mask val loss | no |
| M113 | 1e-5 | 50 | 1 | 1 | mask val loss | no |
| M114 | 1e-3 | 50 | 3 | 1 | mask val loss | no |
| M115 | 1e-4 | 50 | 3 | 1 | mask val loss | no |
| M116 | 1e-5 | 50 | 3 | 1 | mask val loss | no |
| M117 | 1e-3 | 50 | 2 | 1 | mask val loss | no |
| M118 | 1e-4 | 50 | 2 | 1 | mask val loss | no |
| M119 | 1e-5 | 50 | 2 | 1 | mask val loss | no |
| M121 | 1e-3 | 50 | 1 | 1 | val loss | no |
| M122 | 1e-4 | 50 | 1 | 1 | val loss | no |
| M123 | 1e-5 | 50 | 1 | 1 | val loss | no |
| M124 | 1e-3 | 50 | 3 | 1 | val loss | no |
| M125 | 1e-4 | 50 | 3 | 1 | val loss | no |
| M126 | 1e-5 | 50 | 3 | 1 | val loss | no |
| M127 | 1e-3 | 50 | 2 | 1 | val loss | no |
| M128 | 1e-4 | 50 | 2 | 1 | val loss | no |
| M129 | 1e-5 | 50 | 2 | 1 | val loss | no |
| M131 | 1e-3 | 50 | 1 | 10 | val loss | no |
| M132 | 1e-4 | 50 | 1 | 10 | val loss | no |
| M133 | 1e-5 | 50 | 1 | 10 | val loss | no |
| M134 | 1e-3 | 50 | 3 | 10 | val loss | no |
| M135 | 1e-4 | 50 | 3 | 10 | val loss | no |
| M136 | 1e-5 | 50 | 3 | 10 | val loss | no |
| M137 | 1e-3 | 50 | 2 | 10 | val loss | no |
| M138 | 1e-4 | 50 | 2 | 10 | val loss | no |
| M139 | 1e-5 | 50 | 2 | 10 | val loss | no |
| M141 | 1e-3 | 15 | 1 | 10 | val loss | no |
| M142 | 1e-4 | 15 | 1 | 10 | val loss | no |
| M143 | 1e-5 | 15 | 1 | 10 | val loss | no |
| M144 | 1e-3 | 15 | 3 | 10 | val loss | no |
| M145 | 1e-4 | 15 | 3 | 10 | val loss | no |
| M146 | 1e-5 | 15 | 3 | 10 | val loss | no |
| M147 | 1e-3 | 15 | 2 | 10 | val loss | no |
| M148 | 1e-4 | 15 | 2 | 10 | val loss | no |
| M149 | 1e-5 | 15 | 2 | 10 | val loss | no |

Supplementary Table S7: P-values of lsmeans in the statistical analysis of model updating with the factor dataset:model

| Contrast between models | Estimate | SE | df | t ratio | p value |
|---|---|---|---|---|---|
| 10 images COCO vs. 20 images COCO | -6.517 | 0.79 | 944 | -8.294 | $<$0.0001 |
| 10 images COCO vs. 30 images COCO | -6.472 | 0.79 | 944 | -8.237 | $<$0.0001 |
| 10 images COCO vs. 40 images COCO | -10.193 | 0.79 | 944 | -12.973 | $<$0.0001 |
| 10 images COCO vs. 50 images COCO | -12.345 | 0.79 | 944 | -15.712 | $<$0.0001 |
| 10 images COCO vs. 10 images Maize | -17.209 | 0.78 | 944 | -22.015 | $<$0.0001 |
| 10 images COCO vs. 20 images Maize | -16.841 | 0.78 | 944 | -21.519 | $<$0.0001 |
| 10 images COCO vs. 30 images Maize | -18.286 | 0.78 | 944 | -23.366 | $<$0.0001 |
| 10 images COCO vs. 40 images Maize | -18.919 | 0.78 | 944 | -24.174 | $<$0.0001 |
| 10 images COCO vs. 50 images Maize | -18.862 | 0.78 | 944 | -24.060 | $<$0.0001 |
| 20 images COCO vs. 30 images COCO | 0.045 | 0.79 | 944 | 0.058 | $<$0.0001 |
| 20 images COCO vs. 40 images COCO | -3.676 | 0.79 | 944 | -4.679 | $<$0.0001 |
| 20 images COCO vs. 50 images COCO | -5.828 | 0.79 | 944 | -7.417 | $<$0.0001 |
| 20 images COCO vs. 10 images Maize | -10.692 | 0.78 | 944 | -13.678 | $<$0.0001 |
| 20 images COCO vs. 20 images Maize | -10.324 | 0.78 | 944 | -13.192 | $<$0.0001 |
| 20 images COCO vs. 30 images Maize | -11.769 | 0.78 | 944 | -15.039 | $<$0.0001 |
| 20 images COCO vs. 40 images Maize | -12.402 | 0.78 | 944 | -15.847 | $<$0.0001 |
| 20 images COCO vs. 50 images Maize | -12.345 | 0.78 | 944 | -15.748 | $<$0.0001 |
| 30 images COCO vs. 40 images COCO | -3.721 | 0.79 | 944 | -4.736 | $<$0.0001 |
| 30 images COCO vs. 50 images COCO | -5.873 | 0.79 | 944 | -7.475 | $<$0.0001 |
| 30 images COCO vs. 10 images Maize | -10.737 | 0.78 | 944 | -13.736 | $<$0.0001 |
| 30 images COCO vs. 20 images Maize | -10.369 | 0.78 | 944 | -13.250 | $<$0.0001 |
| 30 images COCO vs. 30 images Maize | -11.815 | 0.78 | 944 | -15.097 | $<$0.0001 |
| 30 images COCO vs. 40 images Maize | -12.447 | 0.78 | 944 | -15.905 | $<$0.0001 |
| 30 images COCO vs. 50 images Maize | -12.391 | 0.78 | 944 | -15.805 | $<$0.0001 |
| 40 images COCO vs. 50 images COCO | -2.152 | 0.79 | 944 | -2.739 | $<$0.0001 |
| 40 images COCO vs. 10 images Maize | -7.016 | 0.78 | 944 | -8.976 | $<$0.0001 |
| 40 images COCO vs. 20 images Maize | -6.648 | 0.78 | 944 | -8.495 | $<$0.0001 |
| 40 images COCO vs. 30 images Maize | -8.093 | 0.78 | 944 | -10.342 | $<$0.0001 |
| 40 images COCO vs. 40 images Maize | -8.726 | 0.78 | 944 | -11.150 | $<$0.0001 |
| 40 images COCO vs. 50 images Maize | -8.669 | 0.78 | 944 | -11.059 | $<$0.0001 |
| 50 images COCO vs. 10 images Maize | -4.864 | 0.78 | 944 | -6.223 | $<$0.0001 |
| 50 images COCO vs. 20 images Maize | -4.496 | 0.78 | 944 | -5.745 | 0.0001 |
| 50 images COCO vs. 30 images Maize | -5.942 | 0.78 | 944 | -7.592 | 0.0001 |
| 50 images COCO vs. 40 images Maize | -6.574 | 0.78 | 944 | -8.401 | 0.1598 |
| 50 images COCO vs. 50 images Maize | -6.518 | 0.78 | 944 | -8.314 | 0.1868 |
| 10 images Maize vs. 20 images Maize | 0.368 | 0.78 | 944 | 0.474 | 0.2230 |
| 10 images Maize vs. 30 images Maize | -1.077 | 0.78 | 944 | -1.388 | 0.4552 |
| 10 images Maize vs. 40 images Maize | -1.710 | 0.78 | 944 | -2.203 | 0.5115 |
| 10 images Maize vs. 50 images Maize | -1.653 | 0.78 | 944 | -2.123 | 0.6974 |
| 20 images Maize vs. 30 images Maize | -1.446 | 0.78 | 944 | -1.858 | 0.9309 |
| 20 images Maize vs. 40 images Maize | -2.078 | 0.78 | 944 | -2.671 | 0.9984 |
| 20 images Maize vs. 50 images Maize | -2.021 | 0.78 | 944 | -2.591 | 0.9993 |
| 30 images Maize vs. 40 images Maize | -0.633 | 0.78 | 944 | -0.813 | 1.0000 |
| 30 images Maize vs. 50 images Maize | -0.576 | 0.78 | 944 | -0.738 | 1.0000 |
| 40 images Maize vs. 50 images Maize | 0.057 | 0.78 | 944 | 0.073 | 1.0000 |

Supplementary Table S8: P-values of lsmeans in the statistical analysis of model updating with the factor model:trainingsetsize

| Contrast between models | Estimate | SE | df | t ratio | p value |
|---|---|---|---|---|---|
| 10 images COCO vs. 20 images COCO | -6.52 | 0.79 | 944 | -8.29 | ¡0.0001 |
| 10 images COCO vs. 30 images COCO | -6.47 | 0.79 | 944 | -8.24 | ¡0.0001 |
| 10 images COCO vs. 40 images COCO | -10.19 | 0.79 | 944 | -12.97 | ¡0.0001 |
| 10 images COCO vs. 50 images COCO | -12.34 | 0.79 | 944 | -15.71 | ¡0.0001 |
| 10 images COCO vs. 10 images Maize | -17.21 | 0.78 | 944 | -22.02 | ¡0.0001 |
| 10 images COCO vs. 20 images Maize | -16.84 | 0.78 | 944 | -21.52 | ¡0.0001 |
| 10 images COCO vs. 30 images Maize | -18.29 | 0.78 | 944 | -23.37 | ¡0.0001 |
| 10 images COCO vs. 40 images Maize | -18.92 | 0.78 | 944 | -24.17 | ¡0.0001 |
| 10 images COCO vs. 50 images Maize | -18.86 | 0.78 | 944 | -24.06 | ¡0.0001 |
| 20 images COCO vs. 40 images COCO | -3.68 | 0.79 | 944 | -4.68 | ¡0.0001 |
| 20 images COCO vs. 50 images COCO | -5.83 | 0.79 | 944 | -7.42 | ¡0.0001 |
| 20 images COCO vs. 10 images Maize | -10.69 | 0.78 | 944 | -13.68 | ¡0.0001 |
| 20 images COCO vs. 20 images Maize | -10.32 | 0.78 | 944 | -13.19 | ¡0.0001 |
| 20 images COCO vs. 30 images Maize | -11.77 | 0.78 | 944 | -15.04 | ¡0.0001 |
| 20 images COCO vs. 40 images Maize | -12.4 | 0.78 | 944 | -15.85 | ¡0.0001 |
| 20 images COCO vs. 50 images Maize | -12.35 | 0.78 | 944 | -15.75 | ¡0.0001 |
| 30 images COCO vs. 40 images COCO | -3.72 | 0.79 | 944 | -4.74 | ¡0.0001 |
| 30 images COCO vs. 50 images COCO | -5.87 | 0.79 | 944 | -7.48 | ¡0.0001 |
| 30 images COCO vs. 10 images Maize | -10.74 | 0.78 | 944 | -13.74 | ¡0.0001 |
| 30 images COCO vs. 20 images Maize | -10.37 | 0.78 | 944 | -13.25 | ¡0.0001 |
| 30 images COCO vs. 30 images Maize | -11.81 | 0.78 | 944 | -15.1 | ¡0.0001 |
| 30 images COCO vs. 40 images Maize | -12.45 | 0.78 | 944 | -15.91 | ¡0.0001 |
| 30 images COCO vs. 50 images Maize | -12.39 | 0.78 | 944 | -15.81 | ¡0.0001 |
| 40 images COCO vs. 10 images Maize | -7.02 | 0.78 | 944 | -8.98 | ¡0.0001 |
| 40 images COCO vs. 20 images Maize | -6.65 | 0.78 | 944 | -8.49 | ¡0.0001 |
| 40 images COCO vs. 30 images Maize | -8.09 | 0.78 | 944 | -10.34 | ¡0.0001 |
| 40 images COCO vs. 40 images Maize | -8.73 | 0.78 | 944 | -11.15 | ¡0.0001 |
| 40 images COCO vs. 50 images Maize | -8.67 | 0.78 | 944 | -11.06 | ¡0.0001 |
| 50 images COCO vs. 10 images Maize | -4.86 | 0.78 | 944 | -6.22 | ¡0.0001 |
| 50 images COCO vs. 20 images Maize | -4.5 | 0.78 | 944 | -5.75 | ¡0.0001 |
| 50 images COCO vs. 30 images Maize | -5.94 | 0.78 | 944 | -7.59 | ¡0.0001 |
| 50 images COCO vs. 40 images Maize | -6.57 | 0.78 | 944 | -8.4 | ¡0.0001 |
| 50 images COCO vs. 50 images Maize | -6.52 | 0.78 | 944 | -8.31 | ¡0.0001 |
| 40 images COCO vs. 50 images COCO | -2.15 | 0.79 | 944 | -2.74 | 0.1600 |
| 20 images Maize vs. 40 images Maize | -2.08 | 0.78 | 944 | -2.67 | 0.1900 |
| 20 images Maize vs. 50 images Maize | -2.02 | 0.78 | 944 | -2.59 | 0.2200 |
| 10 images Maize vs. 40 images Maize | -1.71 | 0.78 | 944 | -2.2 | 0.4600 |
| 10 images Maize vs. 50 images Maize | -1.65 | 0.78 | 944 | -2.12 | 0.5100 |
| 20 images Maize vs. 30 images Maize | -1.45 | 0.78 | 944 | -1.86 | 0.7000 |
| 10 images Maize vs. 30 images Maize | -1.08 | 0.78 | 944 | -1.39 | 0.9300 |
| 20 images COCO vs. 30 images COCO | 0.05 | 0.79 | 944 | 0.06 | 1.0000 |
| 10 images Maize vs. 20 images Maize | 0.37 | 0.78 | 944 | 0.47 | 1.0000 |
| 30 images Maize vs. 40 images Maize | -0.63 | 0.78 | 944 | -0.81 | 1.0000 |
| 30 images Maize vs. 50 images Maize | -0.58 | 0.78 | 944 | -0.74 | 1.0000 |
| 40 images Maize vs. 50 images Maize | 0.06 | 0.78 | 944 | 0.07 | 1.0000 |

### Supplementary Figures

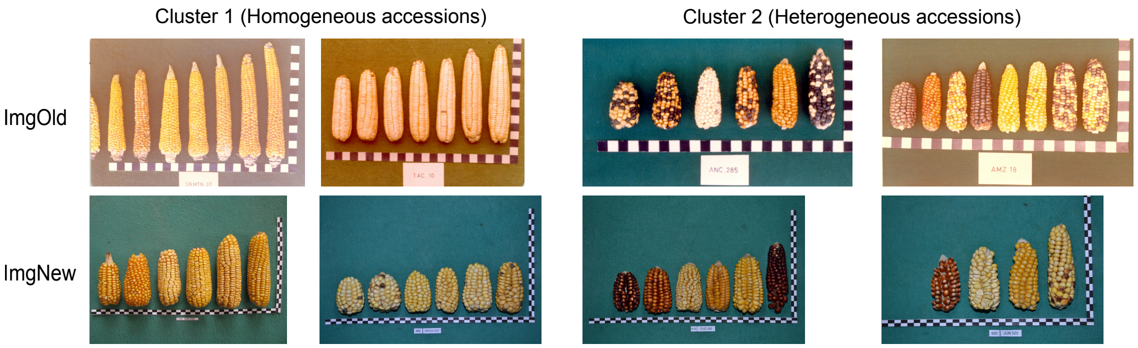


Supplementary Figure S1: Example images of homogenous and hetergenous genebank accessions identified by the multivariate clustering of extracted RGB colors and morphological traits.

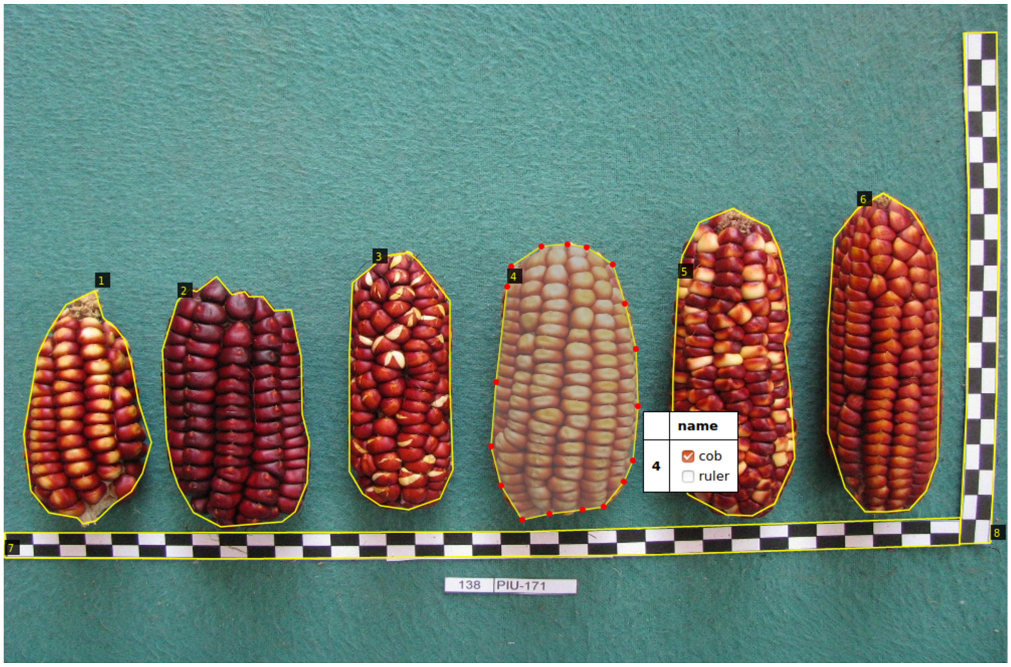


Supplementary Figure S2: Data annotation for *Mask R-CNN* . Polygons were drawn around each object. The ruler was split into the horizontal and the vertical part (to enable also ruler detection on images with only one ruler part) resulting in 2 annotated ruler objects. Finally, the classes of the objects were selected from a checkbox.

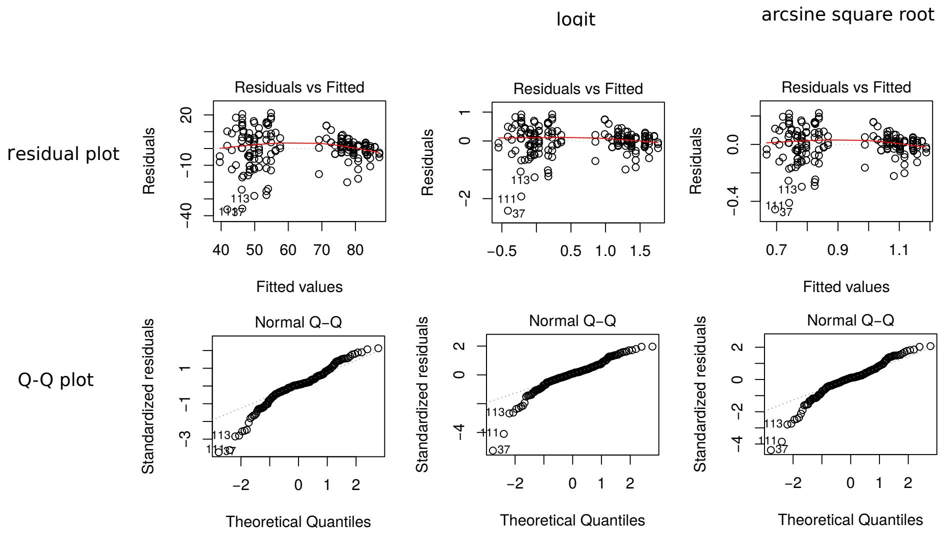


Supplementary Figure S3: Statistical analysis for *Mask R-CNN* parameter selection: Residual plots of different transformations. Logit transformation showed the best results in terms of heterogeneity of variance (residual plot) and normal distribution (QQ-plot) and was therefore applied in the analysis.

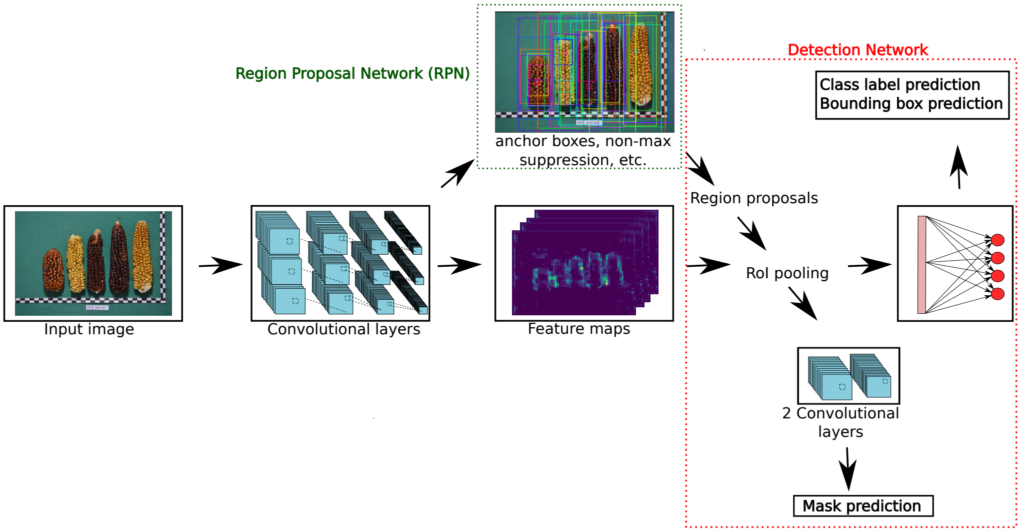


Supplementary Figure S4: Structure of Mask R-CNN visualized with cob images. Mask R-CNN with a block of 2 convolutional layers on top for binary mask prediction.

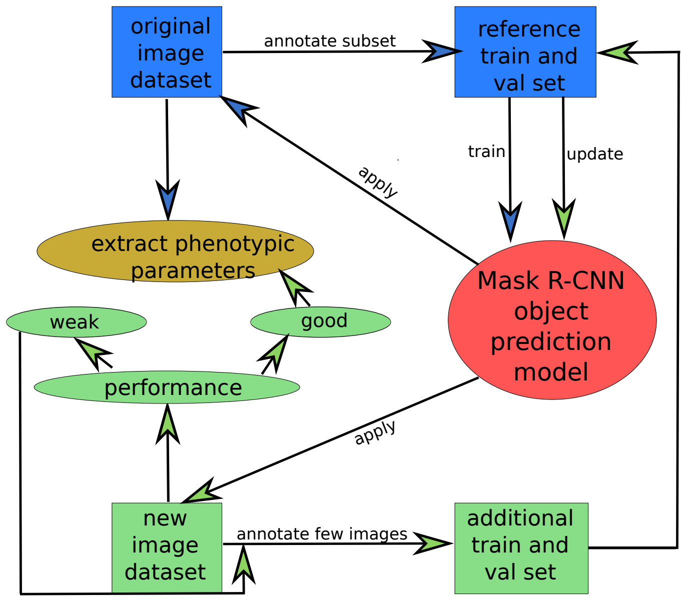


Supplementary Figure S5: Iterative updating scheme of *Mask R-CNN* for application on different datasets and subsequent extraction of phenotypic parameters. In case the model of *Mask R-CNN* does not perform well on a new dataset, the object prediction model is iteratively updated. Two options for the updating step are possible: Either some annotated images from the new dataset are added to the reference train and validation set to update the model (shown here) or the model is re-trained only on these new images.
