## Supplementary Text for "DeepCob: Precise and high-throughput analysis of maize cob geometry using deep learning with an application in genebank phenomics"

### 1 Classical image analysis with the Felzenszwalb-Huttenlocher segmentation

#### 1.1 Implementation of pipeline

We first tested several image processing algorithms from the Python library scikit-image (https://scikit-image.org/) and from the PlantCV library (https://plantcv.danforthcenter.org/) on both ImgOld and ImgNew image datasets. They included the most common image segmentation algoriths, namely Felzenszwalb segmentation, Slic, Quickshift, Sobel operator, Canny edge detector, Otsu thresholding and the Naive Bayes Classifier. Since cobs and ruler differ substantially in color, shape and size, we separately tested and chose the best algorithm for each of them, following the workflow in Figure 1. Additionally, the workflow also required separate adjustments for ImgOld and ImgNew.

For cob detection, the best algorithm, namely Felzenszwalb segmentation, was selected. This image segmentation algorithm [1] has been very successful for other CV tasks due to its graph-based segmentation [2, 3], which segments and image into distinct components by first comparing pixels pairwise and weighting them for reaching maximum homogeneity (lower weights) within a segment, and greatest heterogeneity between segments (higher weights), and accordingly forming edges out of the pixels. The specific parameters had to be fine-tuned separately for the image datasets, to *scale*= 600(ImgOld); 700(ImgNew), *sigma*=0.7(ImgOld); 0.3(ImgNew) and *min_size*=100(ImgOld); 500(ImgNew). A cob detection workflow, returning a list of bounding box coordinates of putative cobs, was set up performing random checks. False positive cob contours were excluded using intervals. ImgOld required an additional color filtering step that removed the blue color channel because of its darker, blue backgrounds.

Ruler detection worked best with the Naive Bayes classifier, where the white and black ruler elements were separated from each other using a mask trained on one sample image. These ruler elements were finally measured using OpenCV for Python, providing intervals for approximate ruler size to sort out false positives. Also, these intervals needed to be set up separately for each image dataset.

In summary, the workflow using traditional image analysis had to be set up separately for ImgOld and ImgNew as well as for the objects cob and ruler, by selecting suitable algorithms and fine-tuning their parameters.

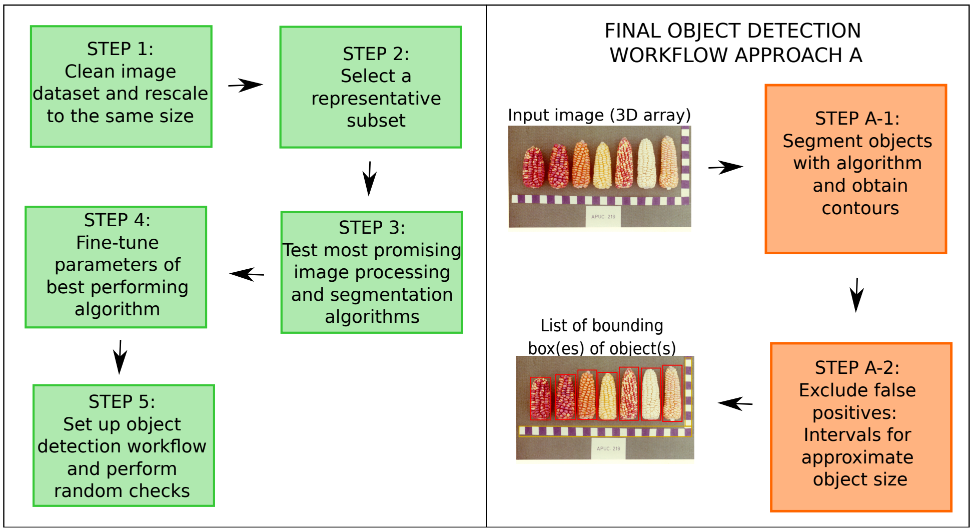


**Figure 1:** Felzenszwalb-Huttenlocher segmentation: Generalized scheme for selecting algorithms, fine-tuning its parameters and setting up a workflow in Traditional Image Analysis (left). The final object detection workflow first segments objects and then excluding false positives (right). This scheme had to be carried out separately for different object classes, like cob and ruler.

#### 1.2 Results of image segmentation with the Felzenszwalb-Huttenlocher segmentation

Of all tested algorithms in Figure2, Felzenszwalb segmentation detected the cobs best and proved to be most robust with respect to the different cob colors, shapes and image brightnesses. The other algorithms performed less well: The Slic segmentation narrowed down the approximate region where all cobs were located, but could not segment individual cobs. Quickshift and Otsu thresholding resulted in blurred masks, in which the cobs are not segmented at all. Canny edge detector outlined the features of yellow maize cobs and ruler elements on a blue and green background quite well, but lead to unsatisfactory results on brown backgrounds and for different colored cobs. The Sobel operator only showed some outlines of ruler elements, but was not at all a reliable segmentation method. Finally, the Bayes Classifier segmented the cobs well from each other in some pictures, but failed for red cobs and darkgreen background. However, this classifier performed best of all algorithms for accurate ruler element segmentation. Based on this comparison, we decided to use Felzenszwalb for classical image segmentation.

Applying Felzenszwalb-Huttenlocher segmentation on the image dataset revealed several critical and reccurring errors in cob detection (Figure3. They included cobs, which were not detected because their color differed from other cobs in the image (Figure3a); cobs that were segmented as group and not individually (Figure3b); and incorrect bounding boxes (Figure3c). In some cases, labels or ruler elements were also segmented as as cobs.

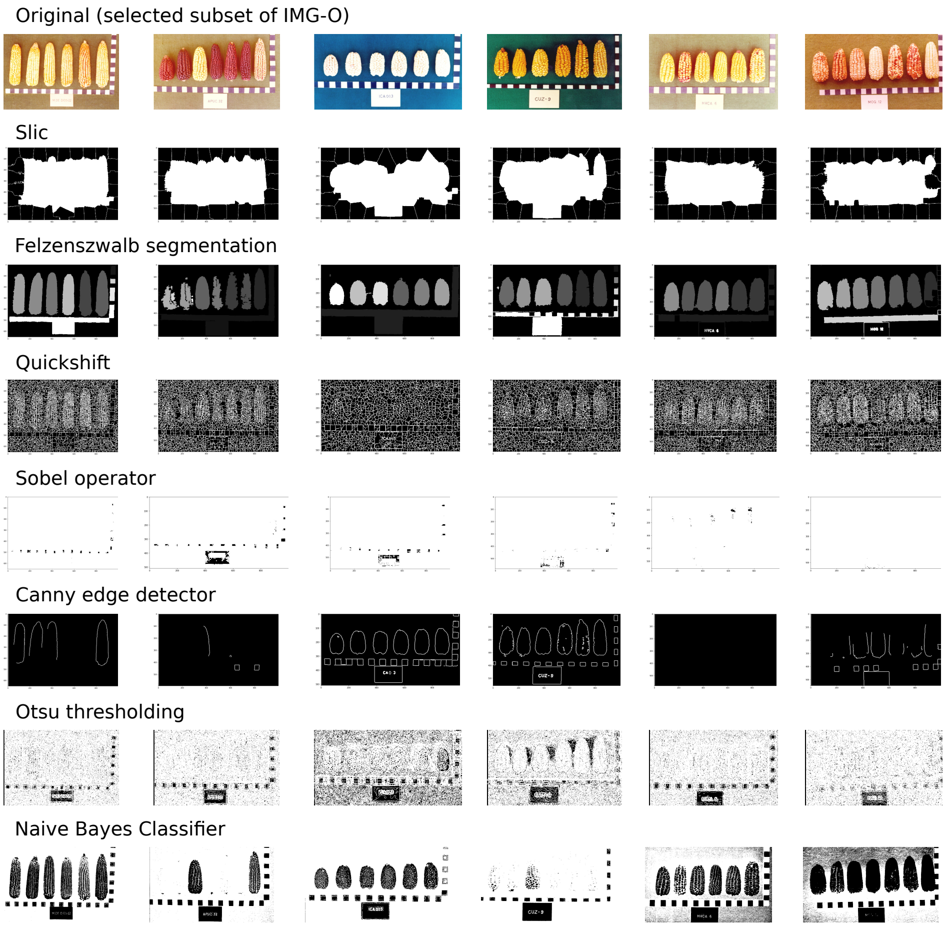


**Figure 2:** Different image processing algorithms presented on a subset of ImgOld. Felzenszwalb segmentation showed the best cob instance segmentation result, more robust to different backgrounds and cob colors than the others.

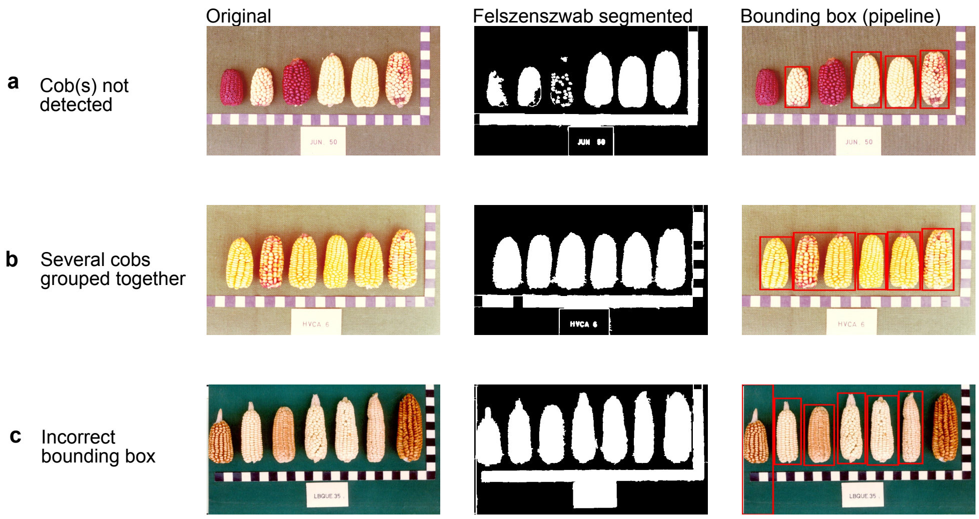


**Figure 3:** Examples of incorrect maize cob detection. (a) Incorrectly detected cobs within an image or no detection of cobs, (b) detection of cobs as a group or (c) incorrect bounding box measures.

Furthermore, despite returning a correct bounding box for most of the cobs, the pixel-wise mask of the cobs (i.e., instance segmentation) was rather inaccurate and did not often show proper cob contours (Figure4). The figure also illustrates another common problem: White and yellow cobs were detected and segmented very well (cob 3 and 7 from the left), whereas the red cob segments often were incomplete and scattered (cob 1,2,4,5,6 from the left), especially on brown backgrounds.

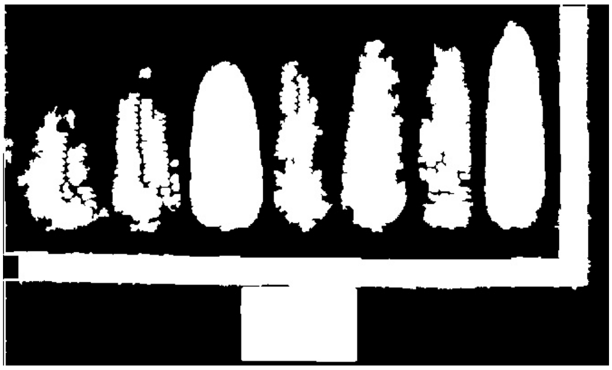

**Figure 4:** Example of an inaccurate cob segmentation result by Felzenszwalb segmentation algorithm which especially fails for red maize cobs.

Overall, success of cob detection was largely dependent of cob color, extent and darkness of cob shadow, background color, and contrast. Due to the multiple issues with classical image segmentation, we used Felzenszwalb-Huttenlocher segmentation mainly as reference for the other two approaches investigated

#### 2 Image segmentation with Window CNN

The complete workflow using a CNN classifier combined with sliding windows is shown in Figure 5. In contrast to the classical image analysis, this approach did not require separate analyses of the ImgOld and ImgNew data sets. Both object classes 'maize cob' and 'ruler' were trained in a single CNN model, although fine-tune for the sliding window process was necessary.

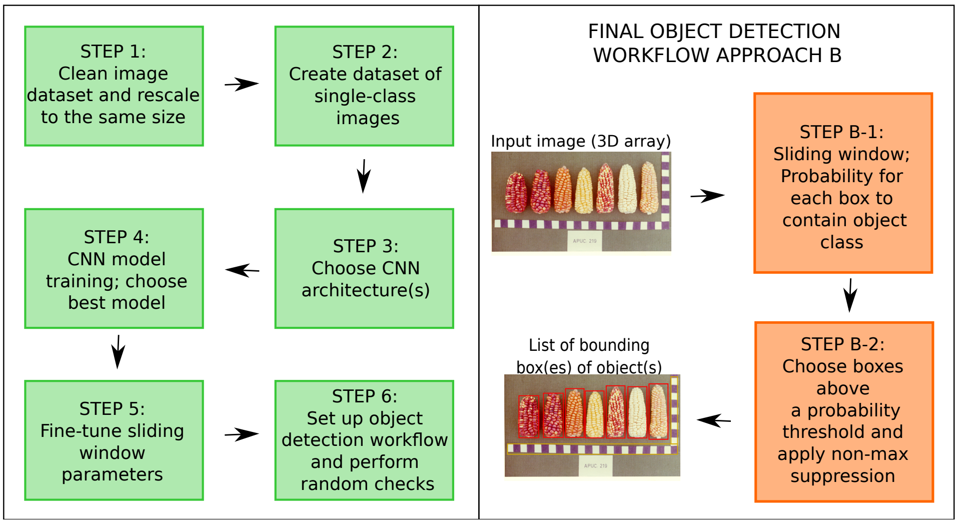


**Figure 5:** The object detection workflow for Window CNN. Left: An image dataset has to be defined to train the CNN classifier. The trained model is then applied together with sliding windows to localize the object. Right: On an input image, a window is slid over the input image, returning each time a probability for the object classes. By applying a certain threshold, prospective bounding boxes are returned after non-max suppression.

The image dataset pre-processing and the CNN model training (Steps 1-4) were carried out only once and contain all different object classes, whereas the following steps had to be carried out separately for each object class. After fine-tuning the sliding window parameters for each class (Step 5), the workflow for the object classes (cob, ruler) differed in these parameters (Step 6).

##### Dataset preparation (Step 1)

First, ImgOld and ImgNew were cleaned, excluding blurred, doubled and not properly named images. Then, the images were re-scaled to 1000x666 pixel. The re-scaling facilitates Step 2 and avoids bias towards single images with a different resolution.

##### Dataset creation for model training (Step 2)

First, a dataset of 37,500 single-class images (B-DATA) was created for training CNN that classify the three classes cob, ruler and background. Usually, a background class is not included in training CNN as simple classifiers, but the highly variable background conditions made this step necessary. To avoid any bias, the image dataset represents a stratified sample of the total available images, having an equal proportion of each class, each image type (ImgOld/ImgNew) and each landrace, as demonstrated in Figure6. Since the CNN requires input images to be of the same size, all images were resized to 80x192 pixel (total of 50GB RAM required), which corresponds best to the mean cob aspect ratio of 2.4. As the manual creation of an image set of this size is very tedious, traditional image processing and cropping techniques as applied with the Felzenszwalb-Huttenlocher segmentation were used to create single-cob images and single images of cob, ruler and background from different image positions. B-DATA was subsequently split into 28,125 training images (75%) and 9,375 validation images (25%).

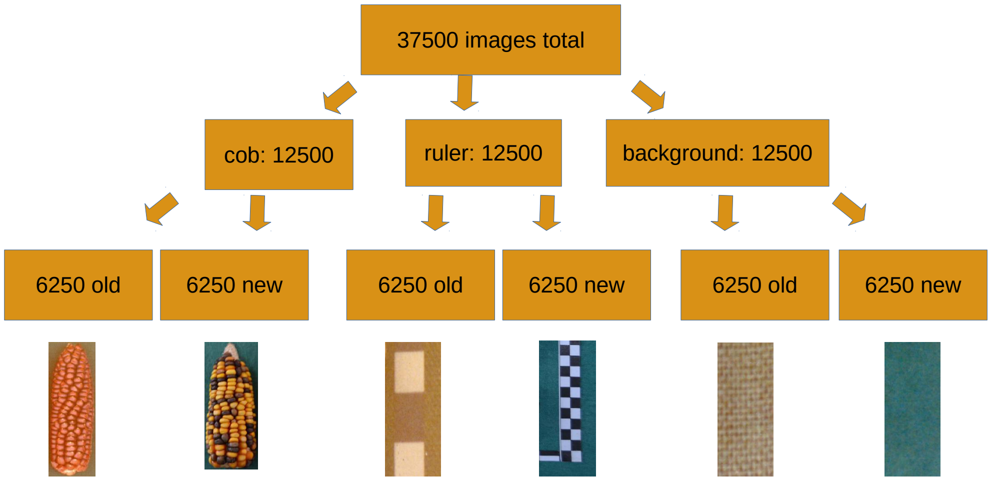


**Figure 6:** Dataset of 37,500 images (B-DATA) created as stratified sample, with equal proportions of each class, image type (ImgOld/ImgNew) and landrace to avoid bias in the training process. All images were resized to 125x300 pixel, which corresponds best to the mean cob aspect ratio of 2.4.

##### Choosing CNN architectures (Step 3)

When combining CNN and sliding windows for object detection, any CNN architecture can be applied in this approach. Two different CNN architectures, differing in complexity, were chosen to investigate the need of complexity to a dataset of only 3 object classes, and trained for classification between cob, ruler and background. Using the typical short notation for net architecture, ShallowNet is a simple CNN with only one convolutional layer and can be written as

$$\text{INPUT} \Rightarrow \text{CONV} \Rightarrow \text{RELU} \Rightarrow \text{FC}$$

After an input layer in which the images of size 80x192 pixel are passed, one single convolutional layer containing 32 filters of size 3x3 is applied and followed by a rectified linear unit (relu) activation. Subsequently, a flattening function decreases the data dimensionality to one dimension, in order to pass it through a FC layer. As there are 3 output classes (cob, ruler, background) in B-DATA, the FC layer has correspondingly 3 output neurons. Overall, giving the image size in B-DATA, this CNN has more than 1 million of trainable parameters (model weights).
Google LeNet [4] is a more complex CNN with 2 convolutional layers, connected by pooling layers and hyperbolic tangent (tanh) activation functions. In short notation, it is represented as

$$\text{INPUT} \Rightarrow \text{CONV} \Rightarrow \text{TANH} \Rightarrow \text{POOL} \Rightarrow \text{CONV} \Rightarrow \text{TANH} \Rightarrow \text{POOL} \Rightarrow \text{FC} \Rightarrow \text{TANH} \Rightarrow \text{FC}$$

.

##### Model training process (Step 4)

A stochastic gradient descent (SGD) optimizer, which minimizes step-wise the training loss to find a global minimum, was used for training the models with a learning rate of 0.0005. The training was executed for 250 epochs, monitoring the training and validation accuracy and loss to identify any signs of over- or underfitting. Finally, the model of the epoch with the lowest validation loss is chosen.

##### Fine-tuning of sliding window parameters (Step 5)

The sliding window parameters a) *Window size* b) *Step size* and c) *Overlap threshold* had to be fine-tuned separately for each object class (cob and ruler).

**Window size** Instead of an image pyramid which scales the window down by keeping a fixed aspect ratio, different windows of different sizes and aspect ratios were fine-tuned to allow a fair chance of all sorts of cobs being correctly localized. The dimensions of these sliding windows are illustrated in Figure7).

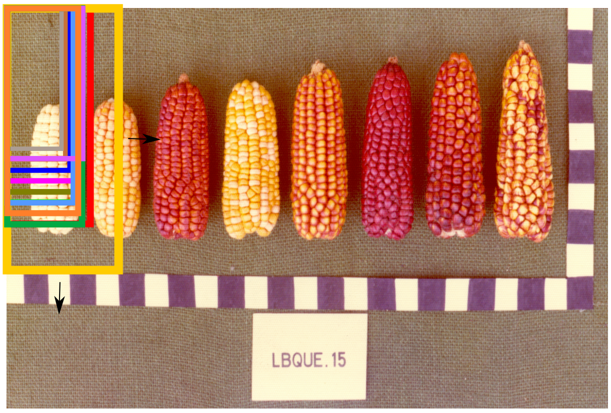


**Figure 7:** Sliding windows of different sizes and aspect ratios that were applied on each image allowing a fair chance to all sorts of cobs to be correctly localized.

**Step size** Different step sizes for the sliding windows of 4,8,12,16 and 20 pixel were tested and finally 16 pixels yielded the best localization, resulting in approximately 10,000 subwindows (all window sizes included) in one image to which the CNN was applied to.

**Overlap threshold** To remove overlapping bounding boxes by keeping only boxes with the highest detection score, post-processing by non-maxima suppression was applied. The best overlapping threshold allowed between two windows was 20%.
Although LeNet showed the highest accuracy, ShallowNet was also tested on these fine-tuned parameters to monitor differences in accurate localization between the models from two different architectures.

##### Setting up workflow and performing random checks (Step 6)

The final object detection workflow was set up with the best-performing model and fine-tuned sliding window parameters to perform it automatically for all given images. After inputting an image as 3D array, a window is slid over the complete image, with the CNN returning the probability for each object class (Step B-1). Here, a strict probability threshold of 0.99999 was set. After applying non-max suppression for excluding overlapping boxes (Step B-2), the final bounding box coordinates for each object are returned.

#### 2.1 Results of Window CNN

In Window CNN a convolutional neural network classifier (CNN) was combined with a sliding window to accurately detect feature bounding boxes in a given image. We first compared two different CNN architectures, ShallowNet and LeNet, with respect to different parameters of the training and detection process. Subsequently, we evaluated the cob and ruler detection performance and its limitations.

##### Accuracy and loss during the CNN training process

Comparing the training process of LeNet and ShallowNet, the process showed no signs of overfitting or underfitting in both cases. The classification loss of LeNet dropped faster and reached its saturation value of nearly 0 faster around epoch 100. ShallowNet approached its slightly higher saturation value of 0.05 slower and it's loss decreased further till epoch 250. These differences can be explained by a more complex net architecture with two convolutional layers of LeNet, allowing for more parameters to be trained during an epoch. In both architectures, the validation loss was always slightly higher than the training loss, which is the expected behavior because the neural net only generalizes from the trained parameters on the validation set, which it has not "seen" before.

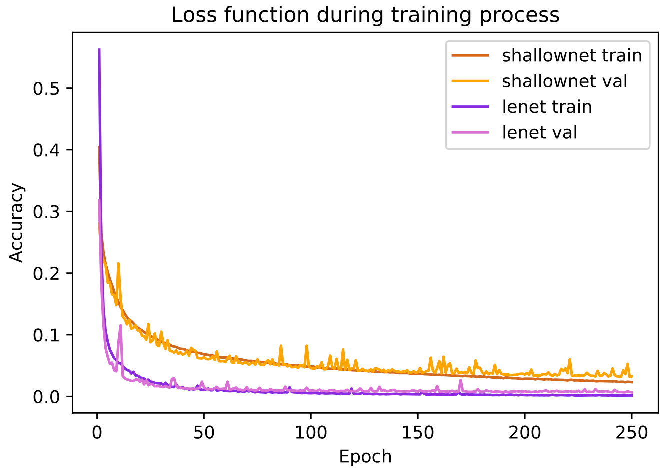


**Figure 8:** Training and validation loss during 250 epochs of training.

**Classification accuracy by class** Regarding the classification accuracy split into the 3 classes (cob, ruler, background) in Table1, ShallowNet and LeNet classified a cob image in the validation set both to 100 % as a cob. Whereas LeNet also classified ruler and background completely with an accuracy of 100%, ShallowNet performed slightly worse with 99% and 98%, respectively. This can be explained by the simpler net architecture, possibly not containing enough parameters to completely classify the ruler and background class, which seem to be more difficult for feature extraction.

**Table 1:** Classification accuracy yielded by the CNN's of ShallowNet and LeNet, split into the 3 classes cob, ruler and background.

| class | ShallowNet | LeNet |
|---|---|---|
| cob | 1.00 | 1.00 |
| ruler | 0.99 | 1.00 |
| background | 0.98 | 1.00 |

Judged by its lower loss and higher classification accuracy, LeNet yielded the best CNN model for the classification dataset that the model was trained on (B-DATA).

##### Cob detection using sliding windows

Testing both net architectures in combination with sliding windows revealed an unexpected behavior. Although the LeNet architecture outperformed ShallowNet in terms of lower loss and higher classification accuracy, in most cases it produces a larger number of boxes than cobs in the image. For example, with the LeNet CNN a cob was often captured by 2 vertical bounding boxes, whereas the ShallowNet model resulted in fewer false positive boxes and was overall more accurate (Figure9).

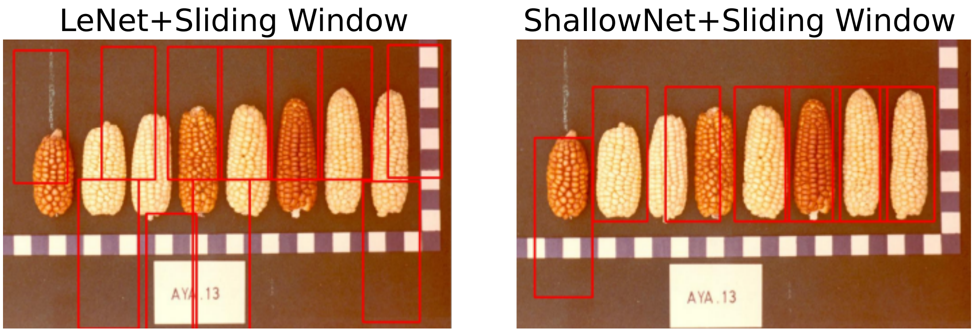


**Figure 9:** Different performance of LeNet and ShallowNet when applied with sliding windows. LeNet showed a higher classification accuracy and a lower loss, but produced many false positives, because cobs were tagged by multiple boxes vertically. In contrast, ShallowNet produced more accurate boxes around the cob.

Although ShallowNet outperformed LeNet in cob localization, it often showed inaccurate bounding boxes around the cobs in situations without high contrast between cob and background, as presented in Figure10. Generally, the boxes sticked to the cob, so the neural net probably learned to detect the features of a cob, but the sliding window approach did not output reliable bounding boxes around cobs.

##### Ruler detection through sliding window

When trying to optimize parameters for ruler detection, like increasing the confidence from 0.9999 to 0.999999 or changing the ROI sizes, the boxes for detecting the rulers were very inaccurate and even spread over regions where no ruler existed at all (Figure11). In addition, some boxes did not only capture rulers, but also included parts of the maize cob. Only in regions with background only, no ruler boxes were present.

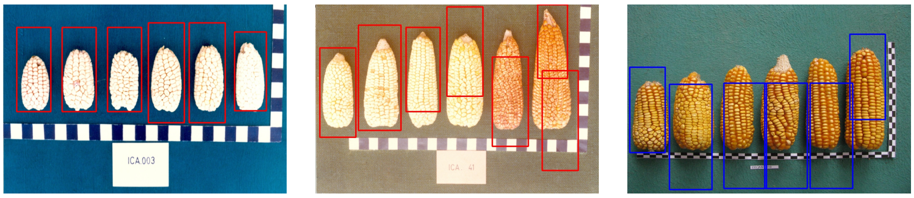


**Figure 10:** Examples for the performance of ShallowNet and sliding window. In images showing high contrast between the cob and the background (left), boxes were very accurate and reliably separated maize cobs from each other. In many other cases, boxes did not capture enclose maize cobs accurately because two boxes were put on the same cob or the box was shifted towards the top or bottom.

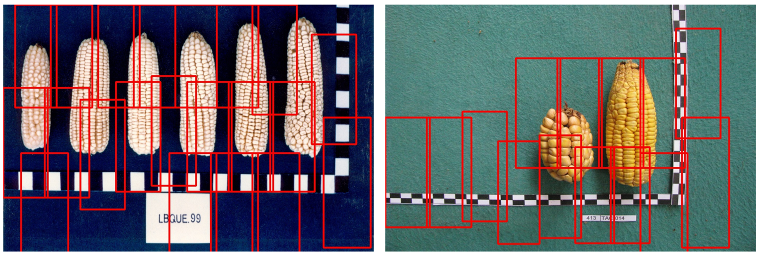


**Figure 11:** Performance of ShallowNet in combination with sliding Window for ruler detection on ImgOld (left) and ImgNew (right). Instead of capturing only the ruler, also boxes containing maize cobs are returned, which rendered ruler detection in Window CNN very unreliable.

### 3 Comparison of all three methods, Felzenszwalb-Huttenlocher segmentation, Window CNN and Mask R-CNN.

#### 3.1 Description of comparison between methods

A comparison for cob detection between all three methods was carried out on a random subsample of 50 images from ImgOld and 50 from ImgNew, which were not included in any of the images used for the taining of the approaches, and the analysis is therefore. The fact that in Approaches A and B cobs were often captured by several boxes and assignment of boxes to an object was not clear did not enable any kind of IoU comparison. Therefore, both a qualitative and a quantitative approach were applied to obtain different kinds of information.

**Qualitative comparison** In the qualitative comparison, several cases of cob detections, as illustrated in Figure12, were compared among the approaches. This comparison counts the occurence of cases, i.e., (1) No detection - where either the cob was not detected or the box deviated largely from the actual position of the cob; (2) A box contains more than one cob; (3) Approximate box - capturing the cob with deviations in its dimensions $<$ approximately 10%; and (4) Accurate box - capturing the cob exactly.

#### 3.2 Results of the comparison

##### Qualitative comparison

In the qualitative comparison, all approaches were compared for their cob detection result according to four defined cases. Table2 highlights the absolute counts of cobs for each case and percentage

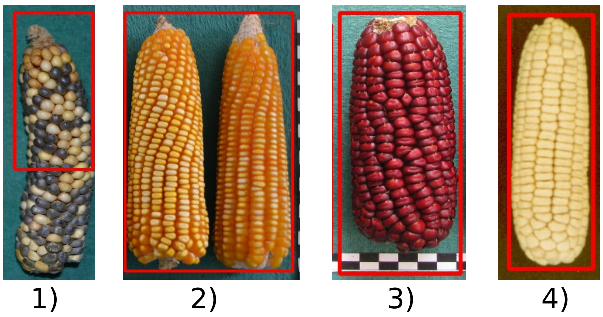

**Figure 12:** Comparison of cob detection between approaches Felzenszwalb-Huttenlocher segmentation, Window CNN, Mask R-CNN for a subset of ImgOld and ImgNew. The cases counted were: 1) No detection, 2) Box containing more than one cob, 3) Approximate box 4) Accurate box.

values, grouped by ImgOld and ImgNew.

**Table 2:** Comparison of cob detection between Felzenszwalb-Huttenlocher segmentation, Window CNN and Mask R-CNN scored for a random subsample of 50 images from ImgOld and 50 images from ImgNew. The values are represented as % of the total cobs in the respective image dataset. The comparison is split into the 4 cases 1) No detection 2) Box> 1 cob 3) Approximate box 4) Accurate box

| | *Felzenszwalb-Huttenlocher segmentation* | *Window CNN* | *Mask R-CNN* |
|---|---|---|---|
| ImgOld (334 cobs total) | | | |
| 1) No detection | 84 (25.1%) | 261 (78.1%) | 0 (0.0%) |
| 2) Box>1Cob | 14 (4.2%) | 1 (0.3%) | 0 (0.0%) |
| 3) Approximate Box | 29 (8.7%) | 71 (21.3%) | 7 (2.1%) |
| 4) Accurate Box | 207 (62.0%) | 1 (0.3%) | 327 (97.9%) |
| ImgNew (235 cobs total) | | | |
| 1) No detection | 10 (4.3%) | 194 (82.6%) | 0 (0.0%) |
| 2) Box>1Cob | 6 (2.6%) | 0 (0%) | 0 (0.0%) |
| 3) Approximate Box | 15 (6.4%) | 39 (15.2%) | 19 (8%) |
| 4) Accurate Box | 204(86.8%) | 2 (0.8%) | 214 (91.2%) |

It reveals that the percentage composition of the four different cases is largely similar in ImgOld and ImgNew. With Felzenszwalb-Huttenlocher segmentation, accurate boxes represent the major part, with some boxes being only approximately correct or containing more than one cob. The method performs much better with the ImgNew than with the ImgOld images. For example, in the latter the number of undetected cobs is much higher. Window CNN does not perform well with both ImgOld and ImgNew images, and it shows a high proportion of undetected maize cobs. Mask R-CNN is outstanding with the highest rate of accurate boxes compared to the other two methods and has only a small proportion of approximate boxes only. With Mask R-CNN, all cobs are detected and there are no boxes that contain more than one cob.

##### 3.2.1 Robustness with diverse image data

We also compared the three approaches with respect to their robustness The final workflows of Felzenszwalb-Huttenlocher segmentation, Window CNN and Mask R-CNN, which were fine-tuned to ImgOld and ImgNew, were tested on the very diverse set of ImgCross. As ImgCross differs from ImgOld and ImgNew substantially, like in image orientation, background color, cob characteristic (also spindles without kernels included) and different labels, it is interesting to evaluate the robustness of each approach on such a different dataset.

A set of representative examples of ImgCross showed differences between the detection performance of the approaches (Figure 13). Felzenszwalb-Huttenlocher segmentation did not detect all cobs and spindles but also it did not return any false positive detections. In contrast, Window CNN detected more cobs and spindles, but also more false positives (red labels) and the bounding boxes were less accurate. None of the two methods was able to detect rotated cobs. Mask R-CNN outperformed Felzenszwalb-Huttenlocher segmentation and Window CNN by detecting all cobs (left) and spindles (right), and at least one rotated cob (middle). It also did not return any false positives, and created bounding boxes and masks of the detected objects with very high accuracy.

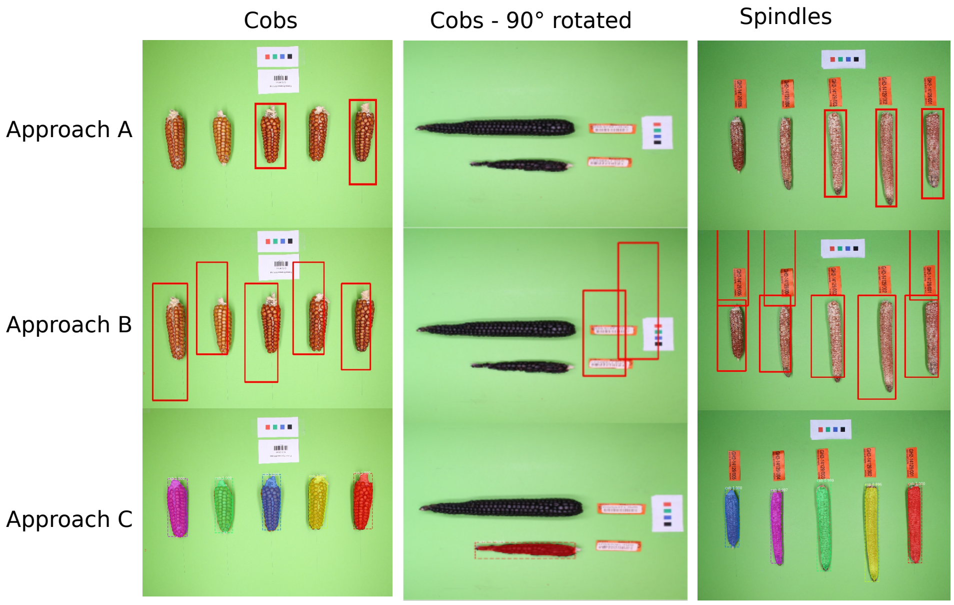


**Figure 13:** Demonstration of the robustness of Felzenszwalb-Huttenlocher segmentation, Window CNN and Mask R-CNN methods on a different image dataset (ImgCross). Mask R-CNN performed in this task and not only detected cobs, but also cob spindles reliably.

We also measured the average script runtime of cob detection on a 16 threads CPU, measured over 100 random images. Runtime was shortest for Felzenszwalb-Huttenlocher segmentation with only 1.2 seconds per image, followed by Mask R-CNN with 3.6 s and Window CNN with 46.7 s. Therefore, Mask R-CNN was more than 12 times faster than Window CNN.
